# Influenza A virus modulation of *Streptococcus pneumoniae* infection using ex vivo transcriptomics in a human primary lung epithelial cell model reveals differential host glycoconjugate uptake and metabolism

**DOI:** 10.1101/2023.01.29.526157

**Authors:** Adonis D’Mello, Jessica R Lane, Jennifer L Tipper, Eriel Martínez, Holly N Roussey, Kevin S Harrod, Carlos J Orihuela, Hervé Tettelin

## Abstract

**Background:** *Streptococcus pneumoniae* (Spn) is typically an asymptomatic colonizer of the nasopharynx but it also causes pneumonia and disseminated disease affecting various host anatomical sites. Transition from colonization to invasive disease is not well understood. Studies have shown that such a transition can occur as result of influenza A virus coinfection.

**Methods:** We investigated the pneumococcal (serotype 19F, strain EF3030) and host transcriptomes with and without influenza A virus (A/California/07 2009 pH1N1) infection at this transition. This was done using primary, differentiated Human Bronchial Epithelial Cells (nHBEC) in a transwell monolayer model at an Air-Liquid Interface (ALI), with multispecies deep RNA-seq.

**Results:** Distinct pneumococcal gene expression profiles were observed in the presence and absence of influenza. Influenza coinfection allowed for significantly greater pneumococcal growth and triggered the differential expression of bacterial genes corresponding to multiple metabolic pathways; in totality suggesting a fundamentally altered bacterial metabolic state and greater nutrient availability when coinfecting with influenza. Surprisingly, nHBEC transcriptomes were only modestly perturbed by infection with EF3030 alone in comparison to that resulting from Influenza A infection or coinfection, which had drastic alterations in thousands of genes. Influenza infected host transcriptomes suggest significant loss of ciliary function in host nHBEC cells.

**Conclusions:** Influenza A virus infection of nHBEC promotes pneumococcal infection. One reason for this is an altered metabolic state by the bacterium, presumably due to host components made available as result of viral infection. Influenza infection had a far greater impact on the host response than did bacterial infection alone, and this included down regulation of genes involved in expressing cilia. We conclude that influenza infection promotes a pneumococcal metabolic shift allowing for transition from colonization to disseminated disease.

**Author summary:** Secondary *Streptococcus pneumoniae* bacterial infections typically occur after influenza A virus respiratory infection. Such coinfections often lead to invasive pneumococcal disease. The mechanisms involved in this process are not well understood. Here, using an *ex vivo* human lung bronchial epithelial cell model, we investigated the biological processes of the host and pneumococcus occurring at this niche, during coinfection with multi-species transcriptomics techniques, and *in vivo* mouse model experimentation. We observed stark differences in global pneumococcal metabolism in different infection states, as well as viral induced epithelial cell changes in ciliary function, potentially aiding pneumococcal dissemination. Overall, this study identified broad and targeted biological processes involved in this host-pathogen interaction.

## Introduction

*Streptococcus pneumoniae* (Spn), a known colonizer of the nasopharynx and a most common cause of community-acquired pneumonia, is also one of the most frequent agents responsible for bacterial coinfection following influenza virus infection (1). Along such lines, considerable historical evidence, including that collected during past influenza pandemics and epidemics, has firmly established that influenza infection starkly predisposes the host to secondary bacterial infections (2). Notably, bacterial coinfections of the respiratory tract alongside a virus are characterized by their greater severity and higher mortality rates when compared to those associated with bacterial pneumonia alone (3, 4).

Multiple studies have investigated the role of influenza and the pneumococcus in superinfections. Preceding influenza infection has been shown to increase pneumococcal burdens in both the upper and lower airway (5). Investigations into the factors that drive host-to-host transmission of Spn revealed increased transmission as result of influenza pre-infection in mice (5), ferrets, and primates. Interestingly, this resulted in reduced influenza transmission (6). Recently, investigators have taken genetic approaches to identify the bacterial genes required for superinfection. For example, using CRISPRi-seq, Liu et al. showed requirement of pneumococcal capsule and the adenylsuccinate synthetase gene purA (7). Rowe et al. identified the bacterial genes required for pneumococcal transmission in a ferret model post influenza A virus (IAV) infection using transposon sequencing (Tn-Seq) (8). Using knockout mutants, they showed that some genes had significant effects on pneumococcal transmission during IAV coinfection, namely, CppA (SP_1449), PiaA (SP_1032), and ComD (SP_2236).

Sequencing-based studies have also contributed to our molecular understanding of events during IAV/Spn coinfection. For example, Reinoso-Vizcaíno et al. showed that the pneumococcal two-component system SirRH increases pneumococcal survival in influenza infected A549 cells, a lung epithelial cell line (9). Pettigrew et al. demonstrated preincubation of influenza with the pneumococcus prior to host infection caused an increase in virulence and bacteremia (10). As for the host’s influence on this process, several major genes and chemokines have been identified as being vital components of the host response. These include IL-17, IL-23, IL-12, CD200, and type I interferon genes (3, 11). Their overexpression is also thought to contribute to the excessive inflammatory response that characterizes secondary bacterial infection and contributes to its severity. These studies, some of which are discussed below in the context of our results, offer conclusive pneumococcal and host markers of superinfection. So far, the bulk of research efforts have been focused on the bacterial factors required for infection and superinfection, and the host response to infection by IAV and Spn. Importantly a comprehensive study on how Spn alters its gene expression and in turn physiology in context of viral infection remains unknown and this is likely to reveal new insights into the basis of disease.

We hypothesized that influenza infection not only skews host defense, rendering the airway more permissive to pneumococcal persistence, but also alters the host environment in manner that alters and promotes bacterial virulence. Here using multispecies RNA-seq we tested this hypothesis by exploring the transcriptome of pneumococcus strain EF3030, a naturally only modestly virulent strain of Spn, during infection of differentiated primary human lung epithelial cells in the presence and absence of influenza A virus pH1N1. Our results shed light on the major genes and pathways involved in the superinfection process from a bacterial and host perspective.

## Results

### *Streptococcus pneumoniae* EF3030 engages in distinct gene expression profiles on primary human lung epithelial cells with and without influenza A pH1N1

To acquire the human lung epithelial and pneumococcal transcriptome in the context of influenza presence or absence, we infected primary nHBEC cells with IAV pH1N1 for 72 hours followed by superinfection with Spn EF3030 for 6h (Figure 1, See Methods). Subsequently, cells were treated with RNA stabilizing reagent, RNA isolated, and multispecies RNA-seq was performed. One caveat of multispecies RNA-seq that commonly occurs is significantly lower sequencing read amounts for the less dominant species. To ensure adequate sequencing depth for robust interrogation of the EF3030 transcriptome, we generated saturation curves for all pneumococcus-infected samples (Supplemental Figure 1A). We observed high coverage for in vitro EF3030 and Host+EF3030+pH1N1 samples, as those saturation curves plateau for over 95% of EF3030 protein coding genes. Host+EF3030 samples were also approaching the same level of saturation for those same genes, giving confidence that our analyses of bacterial signals across all conditions tested are robust.

**Figure 1.**
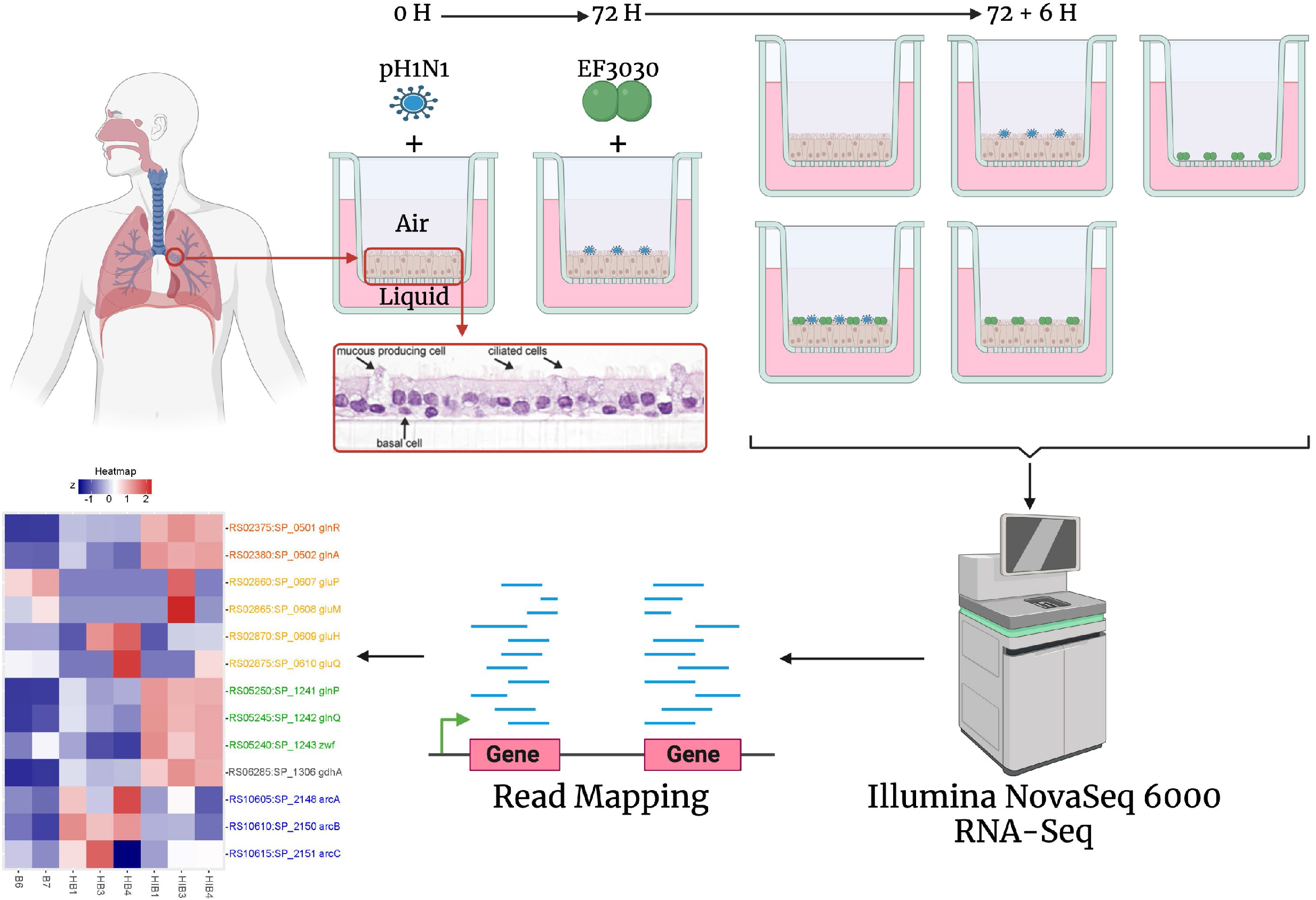
Overview of the study. Airway epithelium was isolated from lung transplant material from a single human donor and purified for cryopreservation. nHBEC cells were thawed and cultured in media for 2 weeks until confluence was reached and transferred to an Air-Liquid Interface system, stimulating differentiation into epithelial phenotypes. nHBEC were then infected or mock infected with pH1N1 virus for 72 hours, followed by *S. pneumoniae* EF3030 infection for 6 hours. Conditions included one coinfection condition with pH1N1 and EF3030, two individual infections of either pH1N1 or EF3030, and two controls of uninfected nHBEC cells and EF3030 cells grown alone in the ALI system. RNA was isolated from experimental replicates of these 5 conditions and sequenced using the Illumina NovaSeq 6000 system, followed by RNA-seq read mapping to the EF3030, pH1N1, and human genomes and analysis, using traditional bulk RNA-seq approaches. Image was created with BioRender.com.

The overall reproducibility of transcripts obtained from biological replicates was evaluated using a principal component analysis (PCA) (Figure 2A). The PCA showed the formation of 3 tight clusters corresponding to the different EF3030 sample types. PC1 captured ∼47% of the overall variation in EF3030 gene expression, with the most difference observed between EF3030 grown in media alone, and EF3030 encountering the human lung epithelium (no influenza: Host+EF3030). However, expression profiles of Host+EF3030+pH1N1 formed a cluster located between in vitro EF3030 and Host+EF3030 along PC1. This suggests that multiple EF3030 genes are in a transitional expression state when compared to in vitro EF3030 and Host+EF3030. PC2 also captured a significant amount of variation at ∼25%, separating the presence of pH1N1 infection of the human lung epithelium for EF3030 genes. This also suggests that certain EF3030 genes might respond to the virus directly or indirectly during the lung epithelial cell infection process. One of the key observations we made from the reads mapping to the EF3030 genome (Supplemental Table 1), was that the proportion of reads that mapped to EF3030 were significantly higher for Host+EF3030+pH1N1 relative to Host+EF3030 (Figure 2B), with ∼30x more bacterial reads mapped from Host+EF3030+pH1N1 samples. A similar observation was made for pH1N1 reads in Host+EF3030+pH1N1 compared to Host+pH1N1, albeit it was not as stark of an increase with ∼0.5x more reads mapping. These results are likely not directly representative of true measures of CFUs of EF3030 in these samples, but we observed a 6x increase in bacterial CFUs from an apical wash of infected transwells in our pilot experiments following identical infection protocols (Supplemental Figure 1B). This correlates with known increases in pneumococcal burden upon influenza infection (5).

**Figure 2.**
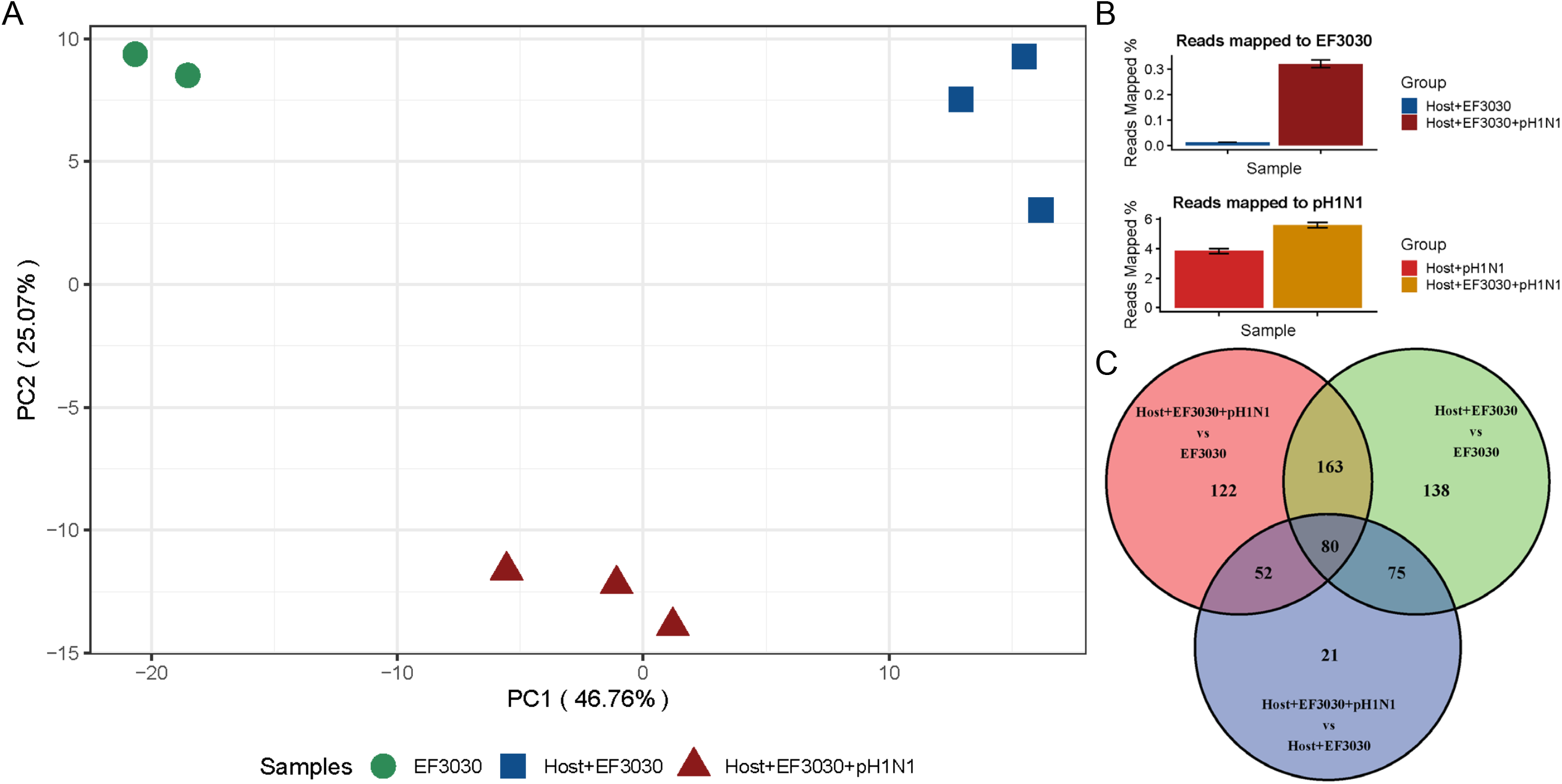
Overview of pneumococcal EF3030 transcriptomes. A) Principal Component Analysis plot of EF3030 gene expression profiles. B) Read mapping percentages to the EF3030 and pH1N1 genomes. C) Venn diagram of genes identified from three differential expression analysis comparisons of EF3030 gene expression.

### Statistical analysis of significantly differentially expressed *Streptococcus pneumoniae* EF3030 genes reveal different responses to primary human lung epithelial cells with and without influenza

To identify the major EF3030 gene responses to epithelial cells, and epithelial cells with pH1N1, we performed a differential expression analysis using DEseq2 (12). We queried differentially expressed (DE) genes for the comparisons of Host+EF3030 relative to in vitro EF3030, Host+EF3030+pH1N1 to in vitro EF3030, and Host+EF3030+pH1N1 to Host+EF3030, considering only DE genes that passed a DEseq2 FDR cutoff of ≤0.05 and an absolute Log_2_ Fold Change (LFC) of ≥1. Figure 1C shows a Venn diagram of the differentially expressed genes for the three comparisons and the intersections of DE genes that were shared among comparisons.

We see that 163 DE genes were shared between the Host+EF3030 vs. EF3030, and Host+EF3030+pH1N1 vs. EF3030 comparisons. This suggests that these genes are modulated by EF3030 upon encounter of the human lung epithelium. There were 80 DE genes in all 3 comparisons, suggesting regulation of these EF3030 genes constitutes a core host-pathogen interaction process. 52 DE genes were also shared between the Host+EF3030+pH1N1 vs. EF3030 and Host+EF3030+pH1N1 vs. Host+EF3030 comparisons, suggesting these EF3030 genes are regulated in the presence of pH1N1. Heatmaps (Supplemental Figure 2) of normalized gene expression values (Supplemental Table 2) of these 3 intersects of DE genes are presented. The heatmaps show a clear differential regulation of the 163 genes, approximately half of which are upregulated on the human lung epithelium regardless of viral infection. A more complex pattern of expression of the 80 genes shared across comparisons. The majority of these show a pattern of strong upregulation of EF3030 genes upon interaction with the human lung epithelium but a relatively lower upregulation in the presence of influenza coinfection. This trend is reminiscent of the sample clustering observed in the PCA plot (Figure 2A) along PC1. For the 52 DE genes, a clear pattern of expression associated with host-pathogen interactions involving the presence of prior viral infection was seen. The Spn EF3030 DE gene lists and the intersections detailed here are provided in Supplemental Table 3.

### Weighted Gene Co-expression Network Analysis (WGCNA) of *Streptococcus pneumoniae* EF3030 transcriptomes captures additional genes associated with host-pathogen interactions

We implemented WGCNA (13) to supplement DE genes following the 3 major patterns of EF3030 gene expression observed in the intersections of Figure 2C. Gene modules were predicted by WGCNA analysis applied to normalized expression values of ∼1900 EF3030 genes (Supplemental Table 1). WGCNA identified 4 modules of EF3030 gene expression, consisting of 835 genes (Figure 3), that fit patterns of interest observed with DE gene sets. The black module consisted of 330 genes regulated specifically when EF3030 encountered the human lung epithelium. The blue module consisted of 193 genes that were more relevant to EF3030 specifically in the absence of a pH1N1 infection, as they were more similar to the in vitro expression profile. The dark green module encompassed 204 genes regulated differently in pH1N1 presence and absence, following the major transitionary pattern of expression seen in the PCA and heatmaps of gene expression (Figure 2A & Supplemental Figure 2, respectively). The final pale violet module comprised 108 genes most relevant to host-pathogen interactions in the presence of pH1N1. In order to visualize the relevance of these modules and assign a numerical value to their importance, we generated PCA plots of the gene subsets from each of these modules in Supplemental Figure 2D-2G. The 330 EF3030 genes of the black module (∼17% of all EF3030 genes) capture ∼78% of the variation along PC1 separating in vitro EF3030 and Host+EF3030, while PC2 only captured subtle differences (∼8%) in biological replicates. The 193 EF3030 genes of the blue module (∼10% of all EF3030 genes) capture ∼84% of the variation along PC1 separating Host+EF3030 interactions specifically in the absence of pH1N1 infection, with PC2 only capturing subtle differences (∼8%) in in vitro EF3030 and Host+EF3030+pH1N1 samples. The 204 EF3030 genes of the green module (∼11% of all EF3030 genes) capture ∼89% of the variation along PC1, revealing genes important to both Host+EF3030 and Host+EF3030+pH1N1 states to different degrees, with PC2 accounting for ∼4% of replicate variation. Lastly, the 108 genes of the pale violet module (∼5.5% of all EF3030 genes) capture ∼88% of the variation along PC1 demonstrating genes most important in the context of prior pH1N1 infection, with PC2 again accounting for ∼4% of replicate variation. These modules encompassing 835 EF3030 genes (∼43% of all EF3030 genes) clearly condense the most condition-regulated genes involved in these specific contexts of host-pathogen interactions and are more associated to specific conditions (pH1N1, Host, or Host+pH1N1) than other genes in this dataset. However, not all the genes identified in modules are necessarily of relevance to the condition considered, and some spurious correlations may exist. Despite this, given that the expression of this subset of the EF3030 pneumococcal transcriptome is highly correlated with Host and pH1N1 presence or absence in multiple replicates, and the high percent variation (78-88%) captured by each module with its respective condition, it is likely that other pneumococci possessing these genes would respond similarly to these host and pH1N1 conditions. The list of all these genes and their normalized expression values are listed in Supplemental Tables 1 & 3.

**Figure 3.**
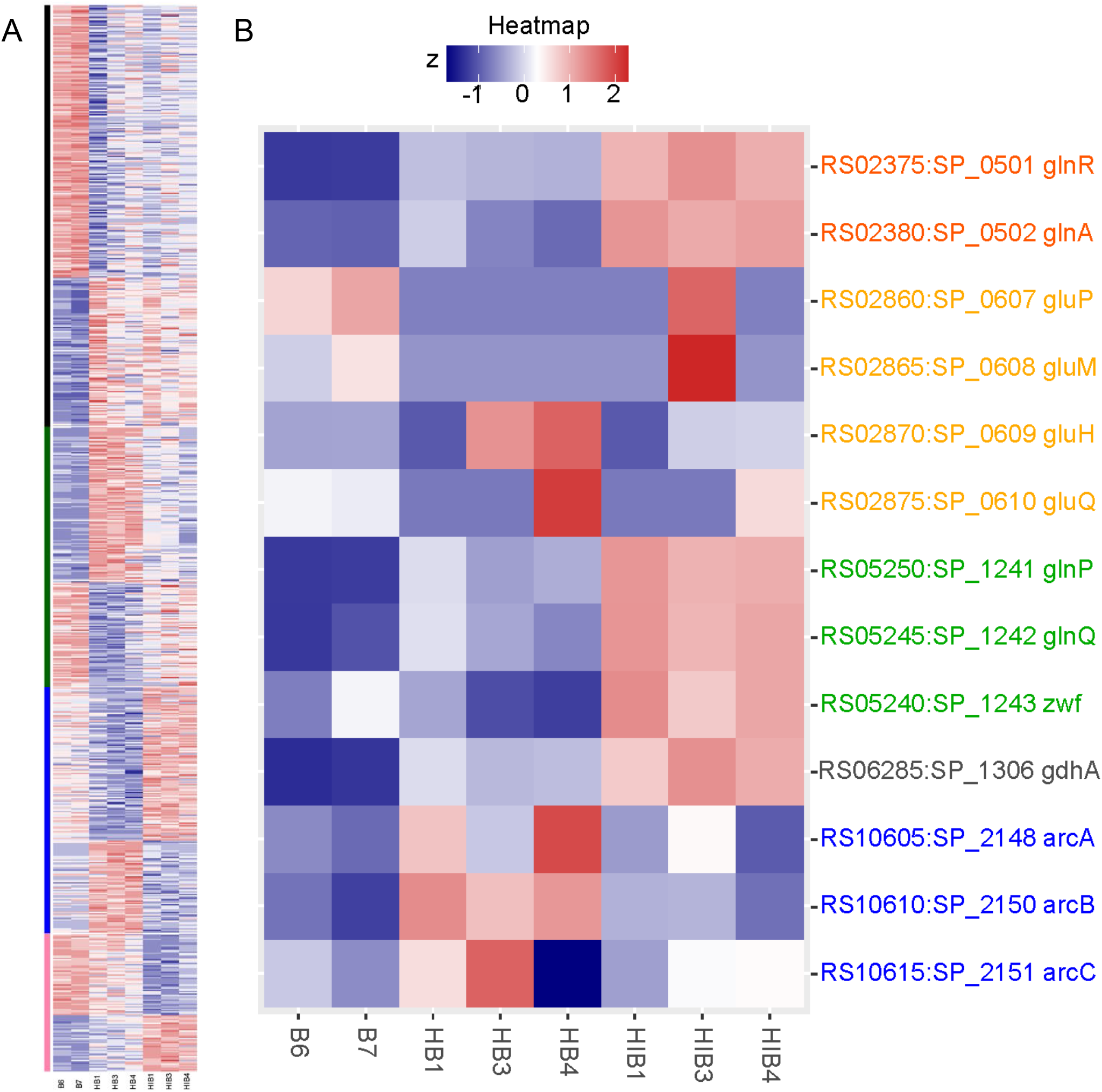
Condition specific pneumococcal EF3030 gene expression patterns. A) Z-scored WGCNA heatmap of normalized expression levels of 835 EF3030 genes clustering in four modules – black, blue, green & pink, from top to bottom on left. B) Z-scored heatmap of expression levels of the EF3030 GlnR regulon, where colored text represents an operon or gene.

As we had hundreds of genes across all 4 modules, we chose to use the eggNOG-mapper (14) to classify the genes in each module, to identify overarching biological processes. The top 3 Clusters of Orthologous Genes (COG) categories in the black module (human lung epithelium response) were: function unknown (COG category S, ∼18% of genes in the module); translation, ribosomal structure and biogenesis (J, ∼13%); and nucleotide transport and metabolism (F, ∼10%). Similarly, for the blue module (EF3030 infection only), categories were: function unknown (S, ∼18%); nucleotide transport and metabolism (F, ∼10%); and carbohydrate transport and metabolism (G, 10%). For the green module (condition-specific gene expression): function unknown (S, ∼17%); carbohydrate transport and metabolism (G, 15%); and inorganic ion transport and metabolism (P, ∼11%). Lastly, for the pale violet module (pH1N1 presence in infection): nucleotide transport and metabolism (F, ∼19%); function unknown (S, ∼16%); and lipid transport and metabolism (I, ∼8%). Complete eggNOG-mapper results and module gene lists are available in Supplemental Table 3.

Despite incomplete annotations of all genes by eggNOG, the ones that were assigned a category represented large subsections of each module. Notably the most affected genes were those involved in diverse aspects of bacterial metabolism. As we were mainly interested in genes in the green and pale violet modules for biological relevance, we further investigated metabolic regulons using the TIGR4 RegPrecise database (15). For instance, we saw multiple genes from the pneumococcal GlnR regulon were DE and found in the pale violet module (pH1N1 presence in infection). A heatmap of the GlnR regulon (Figure 3B) shows that specific genes of the regulon are upregulated during EF3030+pH1N1 infection. In vivo pneumococcal knock out mutant experiments of this regulon will be described in the next section.

Besides this, several operons and genes show specific responses to one of the three conditions we studied, namely, EF3030 in vitro, Host+EF3030 & Host+EF3030+pH1N1. For example, several carbohydrate processing genes such as mannose (EF3030_RS01420 - EF3030_RS01430), galactose (EF3030_RS03030 - EF3030_RS03040), maltose (EF3030_RS10400 - EF3030_RS10410), galactosamine (EF3030_RS00330 - EF3030_RS00350) and sialic acid (EF3030_RS09720 - EF3030_RS09740) are upregulated specifically during EF3030 infection but not during EF3030+pH1N1 coinfection. Whereas pyruvate usage (EF3030_RS04270 - EF3030_RS04275), glycerol (EF3030_RS10795 - EF3030_RS10805), maltodextrin (EF3030_RS10390 - EF3030_RS10395) and ATP synthetase genes (EF3030_RS07065 - EF3030_RS07100) and upregulated in the pH1N1 coinfection state relative to EF3030 infection. Similarly, multiple genes that trend toward upregulation in a pH1N1 coinfection state consisted of utilization of nitrogen sources such as glutamate (EF3030_RS06285, part of the GlnR regulon, Figure 3B), threonine (EF3030_RS02175 - EF3030_RS02200), aspartate (EF3030_RS04905 - EF3030_RS04910) and potentially other branched chain amino acids (EF3030_RS04030). These observations were expanded upon using the manually curated regulons in the Spn TIGR4 RegPrecise database (15). A Z score heatmap of EF3030 gene expression was generated for each regulon that had at least 2 genes differentially expressed. This identified 23 regulons, 14 of which had stark patterns of gene expression and were compiled in Figure 4. The remaining 9 are also displayed in Supplemental Figure 3. Apart from this, a heatmap of known pneumococcal virulence genes (16) was also supplemented into Figure 4.

**Figure 4.**
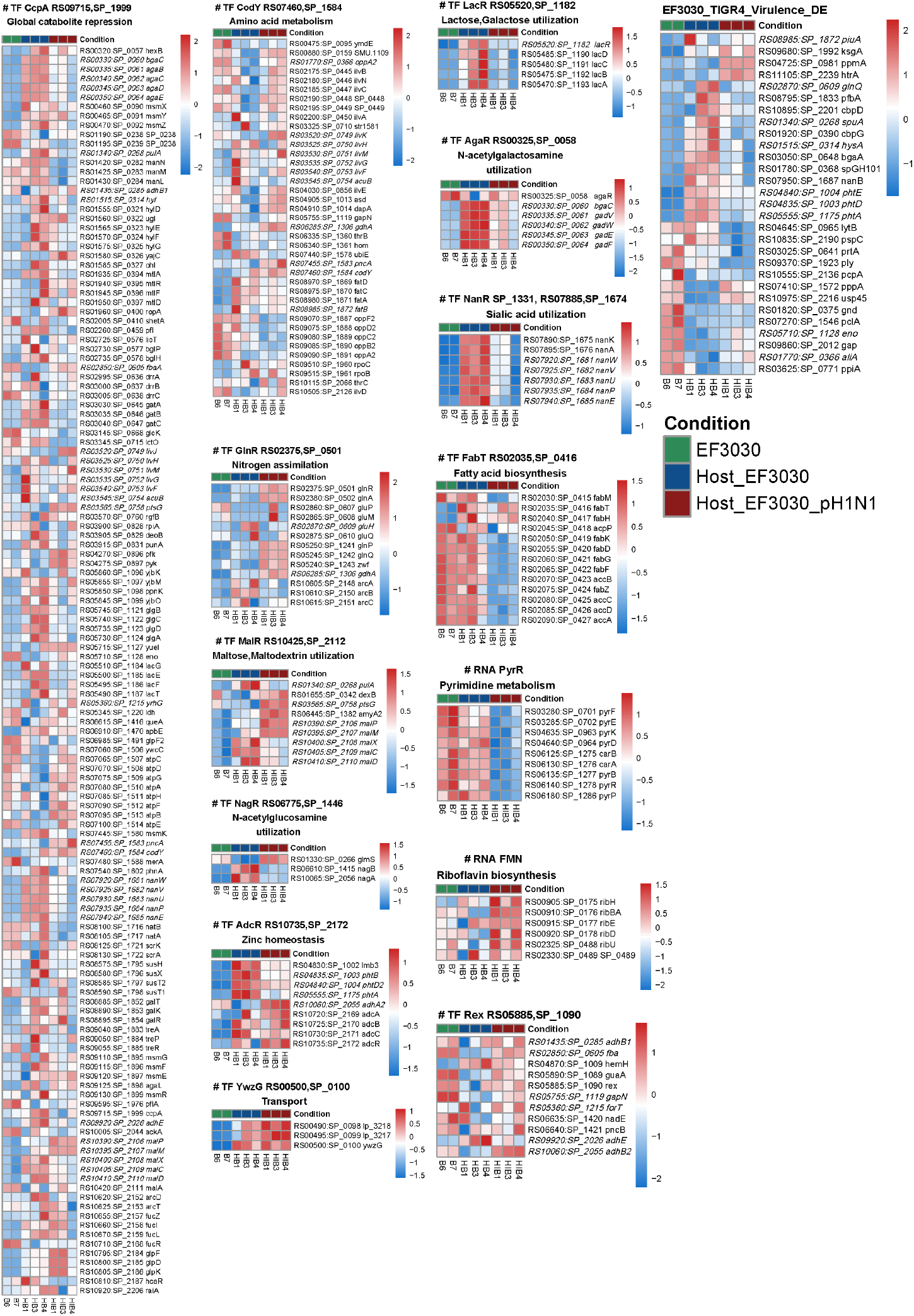
Gene expression profiles of differentially regulated pneumococcal metabolic regulons. Z-scored heatmaps of Regprecise regulons for *S. pneumoniae* strain TIGR4 are presented with strain EF3030 orthologs. Only genes present in both EF3030 and TIGR4 are displayed for each regulon. Genes are ordered by locus tag identifiers, except for the Virulence heatmap which is clustered on gene expression profiles. Italicized fonts indicate genes present in multiple (2-3) regulons. Complete regulon gene lists are available in Supplemental Table 5.

Importantly, we also performed an independent validation of our EF3030 RNA-seq data (the limiting species in our multispecies assay) using the Nanostring platform on the same RNA samples we analyzed. We designed a Nanostring panel of 63 pneumococcal core or near-core pneumococcal genes (Supplemental Table 3) and correlated their observed expression in RNA-seq and Nanostring (Supplemental Figure 4). We observed strong correlations for these 63 genes between the two datasets with spearman correlation (R) values ranging from 0.81 - 0.9. This gave us confidence in our analyses and the conclusions we drew from our EF3030 RNA-seq analyses.

### Influenza A virus infection overwhelmingly alters the human lung epithelium transcriptome

We performed host transcriptome analyses following some of the same approaches as described for the bacterial transcriptome. Host conditions included: control uninfected human lung epithelial cells (Host), Host+EF3030, Host+pH1N1, and Host+EF3030+pH1N1. PCA analysis (Figure 5A) revealed a stark separation along PC1 (∼85% of the variation) between Host+pH1N1 and Host+EF3030+pH1N1 on the right, and Host and Host+EF3030 on the left. The next component PC2 captured only ∼5% of variation between biological replicates. Host and Host+EF3030 samples clustered together with almost no difference, suggesting no major influence on the host as a result of only EF3030 infection (6h) of human lung epithelial cells. Similarly, within pH1N1 containing samples, no clear separation was seen between Host+pH1N1 and Host+EF3030+pH1N1 suggesting that the presence of EF3030 coinfection had no broad effect during pH1N1 infection, potentially due to an overwhelming response to just pH1N1. However, the fact that bacterial gene transcriptomes showed significant differences between EF3030, Host+EF3030 and Host+EF3030+pH1N1, imply that intermixing of biological replicates from a host perspective was not due to bad sample replication but merely the lack of a detectable response to EF3030 during human epithelial lung cell infection and during pH1N1 coinfection in this experimental design.

**Figure 5.**
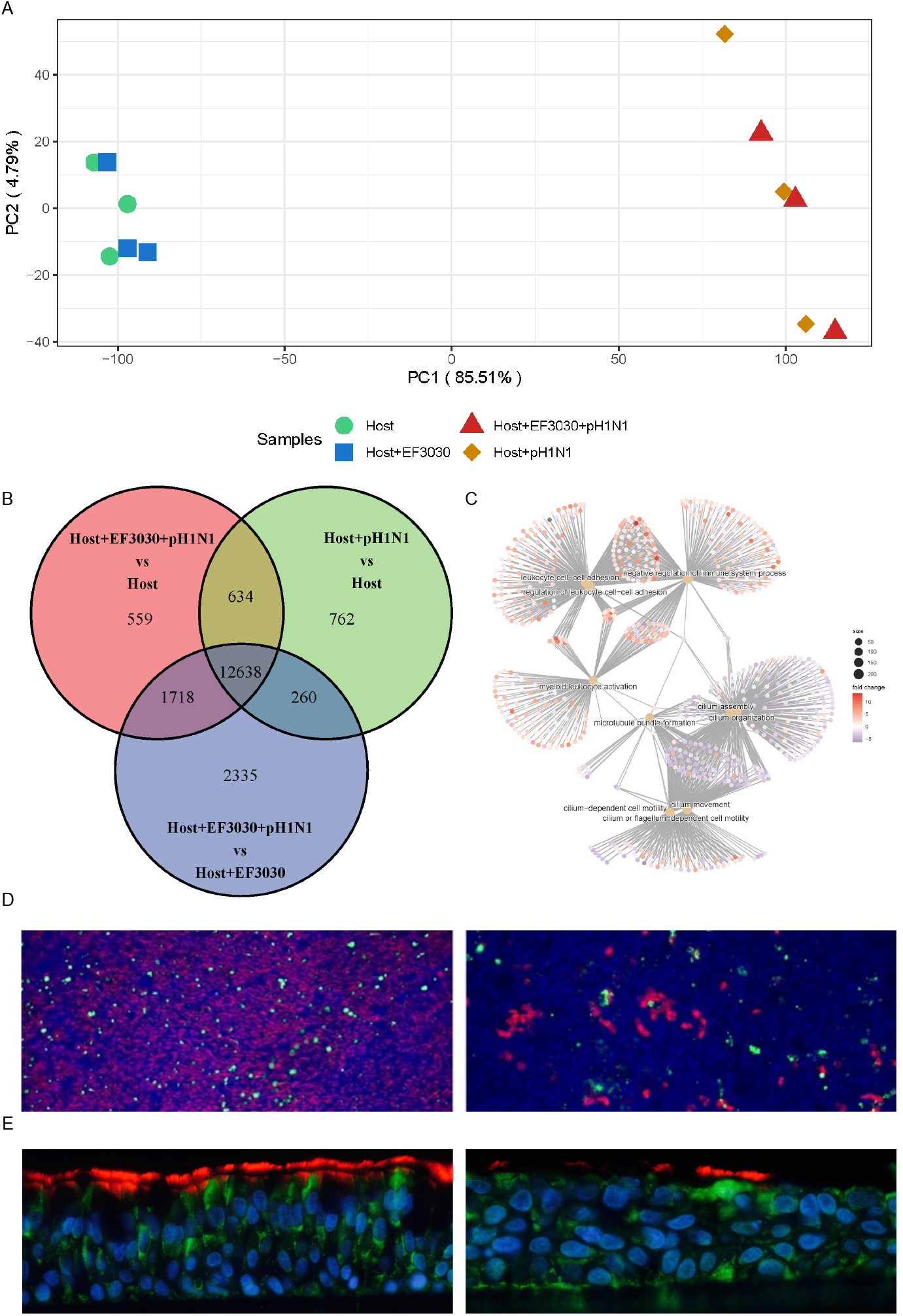
Overview of human nHBEC transcriptomes. A) Principal Component Analysis plot of nHBEC gene expression profiles. B) Venn diagram of genes identified from three differential expression analysis comparisons of nHBEC gene expression. C) Top 10 enriched Gene Ontology biological processes visualized as a gene-concept network plot: inner beige circles represent biological processes and outer circles represent genes colored by Log_2_ Fold Change for the Host+EF3030+pH1N1 vs. Host+EF3030 comparison. D) Immunofluorescence of ALI cultured nHBEC cells: uninfected cells (left); cells infected with pH1N1 for 72 hours (right). Red stain represents cilia and green stain represents mucin (MUC5AC) producing goblet cells, at 20x magnification. E) Immunofluorescence images of ALI from slices of embedded transwells, under the same respective conditions as panel C (cilia in red and CFTR in green).

Using DESeq2 (12), we were able to identify 12638 genes specific to pH1N1 infection of the human lung epithelium, with 2335, 762 and 559 DE genes unique to the comparisons of Host+pH1N1 vs. Host, Host+EF3030+pH1N1 vs. Host, and Host+EF3030+pH1N1 vs. Host+EF3030, respectively (Venn diagram, Figure 5B). Only a few genes (<3) were DE passed an FDR cutoff of ≤0.05 and an absolute Log_2_ Fold Change (LFC) of ≥1 for the comparisons of Host+EF3030 vs. Host, and Host+EF3030+pH1N1 vs. Host+pH1N1. The list of all Host DE genes and subsets are presented in Supplemental Table 4. For the comparisons of Host+pH1N1 vs. Host, Host+EF3030+pH1N1 vs. Host, and Host+EF3030+pH1N1 vs. Host+EF3030, several thousands of genes were DE pass cutoffs, nearly a quarter of all annotated human genes. Most noticeably, 1,000 more genes were DE in Host+EF3030+pH1N1 vs. Host relative to Host+pH1N1 vs. Host, and another 1,500 genes in Host+EF3030+pH1N1 vs. Host+EF3030 when compared to Host+EF3030+pH1N1 vs. Host. This suggests that subsets of genes might be altered to different magnitudes due to the presence of just EF3030 during infection, and some might be due to the presence of just pH1N1 in the context of coinfection. However, it is important to note that because no genes were DE for Host+EF3030 vs. Host, and Host+EF3030+pH1N1 vs. Host+pH1N1, these additional DE gene subsets, though likely modulated during coinfection, may also be DE due to altered responses to EF3030 but not significant enough to meet DE cutoffs in the Host+EF3030 vs. Host, and Host+EF3030+pH1N1 vs. Host+pH1N1 comparisons.

As several thousands of DE genes were identified during infection/coinfection, typical pathway analyses would be overwhelmed with the large numbers of genes as they are optimized to work with much smaller datasets, with the typical upper limit of ∼3,000 DE genes. Hence, we attempted to identify the broader biological processes involved during both pH1N1 infection and EF3030+pH1N1 coinfection using GO term analysis using the R package ClusterProfiler V4.0 (17). Associated GO processes for each comparison are listed in Supplemental Table 4. We saw enrichment of several of the same GO biological processes in all comparisons involving pH1N1 presence vs. pH1N1 absence, mainly involving the upregulation of several cytokine and immune signaling processes, but also a severe decline in cilial regulation, cilial beating and microtubule pathways (Figure 5C). The cilial downregulation was also confirmed phenotypically by the absence of cilia using fluorescence microscopy (Figure 5D).

### Requirement of pneumococcal GlnR regulon associated operons in a mouse lung model of infection

To experimentally validate our RNA-seq results and confirm the biological relevance of some of the differentially regulated host pathways and pneumococcal regulons, we used an in vivo mouse model of infection (Figure 6). We generated 5 bacterial whole operon or gene knockouts of the parts of the GlnR regulon (Figure 3B) in serotype 4 strain TIGR4. We sought to validate our results using EF3030, however preliminary CFU counts of EF3030 lung infections showed low bacterial burdens (data not shown). TIGR4 is more virulent than EF3030 and capable of causing disseminated disease (18). Hence, each of these 5 isogenic mutants, namely ΔgluQHMP, ΔglnPQ/zwf, ΔarcABC, ΔglnRA, and ΔgdhA, were then used in a competition assay between wildtype TIGR4 and mutant, with a 1:1 ratio of knockout to wildtype (Figure 6A). Based on the pattern of gene expression of the GlnR regulon observed in these conditions (Figure 3B), we expected that the ΔglnPQ/zwf, ΔglnRA, and ΔgdhA mutants would be further attenuated in a coinfection with pH1N1, relative to just EF3030 infection. Also, due to the inconsistency of gene expression of ΔgluQHMP and ΔarcABC across conditions and replicates (Figure 3B), we anticipated that these mutants would have no effect and serve as positive controls of the RNA-seq data. In contrast to these predicted results, we observed that only ΔglnPQ/zwf and ΔglnRA were attenuated versus control, and in context of pH1N1 coinfection, ΔglnRA was actually less attenuated rather than hyper-attenuated. However, it is important to note that of all the mutants, the ΔglnRA mutant had a fitness defect indicated in its growth curve (Supplemental Figure 5). Overall, these data indicate that genes with altered expression are not necessarily required, but also that influenza preinfection provides an environment where an otherwise attenuated version of Spn may be able to cause disease.

**Figure 6.**
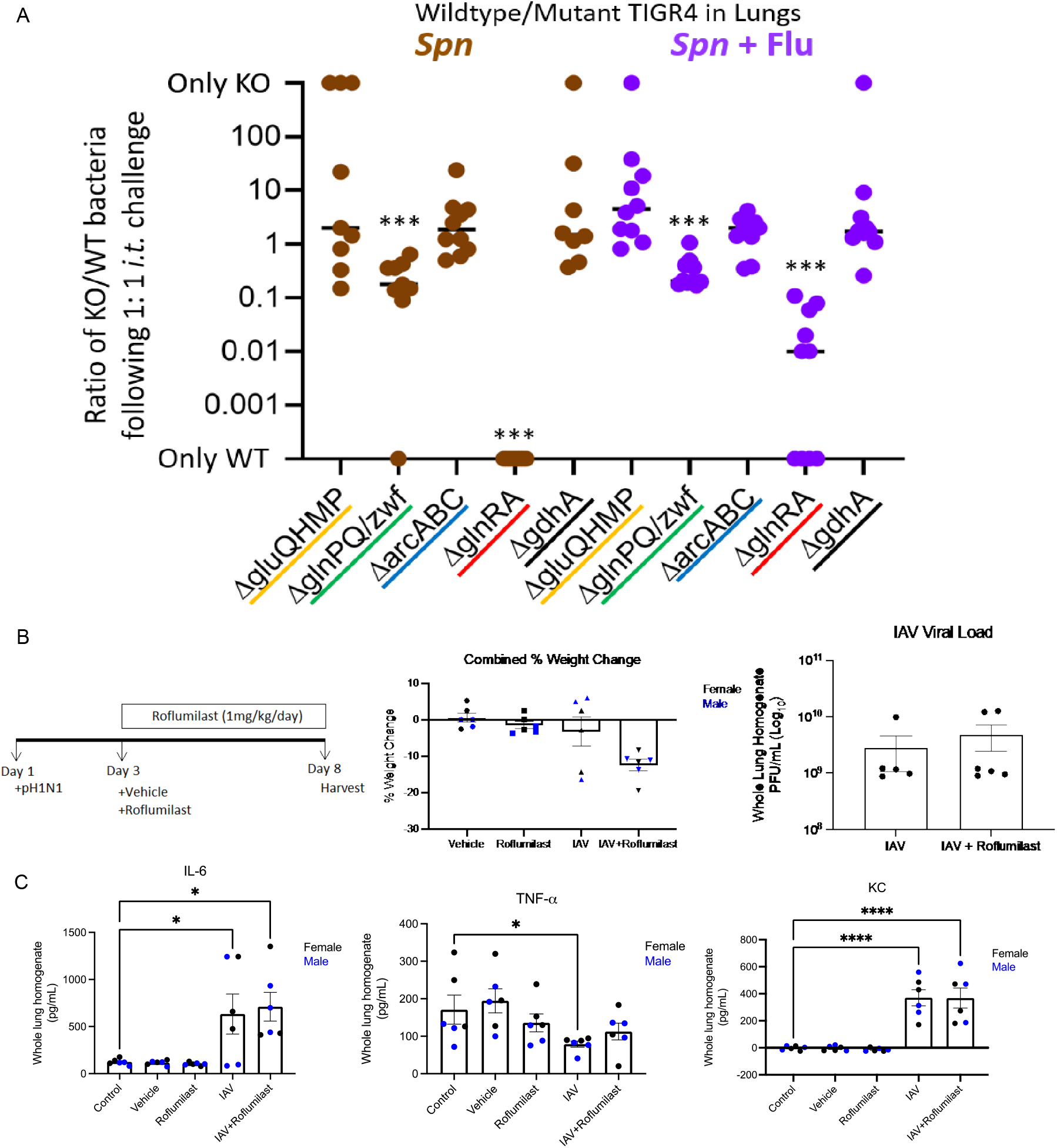
In vivo mouse infection experiments. A) In vivo 1:1 competition experiments between wildtype (WT) TIGR4 and 5 isogenic mutants in 1 gene and 4 operons of the GlnR regulon. Colored lines for the gene and 4 operons correspond to colors in Figure 3B. B) Overview of Roflumilast treatment: experiment design (left), percentage of weight change for infected and Roflumilast treated mice on day 8 post infection (center), and pH1N1 viral burdens in infected and Roflumilast treated mice on day 8 post infection (right). C) ELISA of chemokines IL-6, TNF-α, KC (CXCL8) for infected and Roflumilast treated mice on day 8 post infection. N=6 mice per group and statistical analysis shown is standard error of the mean (SEM). Outliers, if any, were removed. * = p ≤ 0.05; **** = p ≤ 0.0001.

### Roflumilast treatment of IAV infection mice

As for the host response, since it was primarily only due to pH1N1 infection, we focused on pH1N1 regulated GO biological processes, specifically the downregulation of cilial and microtubule proteins (Figure 5) as we had observed it transcriptionally and phenotypically. We hypothesized that pH1N1 infection drives the loss of cilia which in turn affects surface clearance of influenza and any other pneumococcus present during coinfection, ultimately allowing for increased infection of the epithelium by the pathogens. It has been shown that cilia are regulated through the cAMP signaling and the host PDE4 gene acts as an inhibitor (19). We observed that the PDE4 gene is ∼16-fold upregulated in the presence of pH1N1, based on our RNA-seq data. Roflumilast is an FDA approved therapeutic used primarily in COPD (20) that inhibits the PDE4 gene and is a potential avenue for interference of the cilial system, which has shown promise as a potential treatment for RSV infection (21). We hypothesized that the use of Roflumilast to inhibit cilial function in vivo in mice, in the context pH1N1 infection, would enhance mice survivability. As such, we attempted to replicate the experiments performed by Mata et al. (21), as they had demonstrated a significant reduction in viral (RSV and pH1N1) burdens upon roflumilast treatment in female mice. The experimental design was 72 hours of pH1N1 infection of mice followed by oral gavage treatment using Roflumilast at 1mg/kg/day for 5 days (Figure 6B left). However, in contrast to the study by Mata et al. (21), we observed significant weight loss in the infected and roflumilast treated mice (Figure 6B center), and no significant reduction in viral loads after Roflumilast treatment (Figure 6B right). There was also no significant effect of Roflumilast treatment observed in ELISA experiments testing the expression of chemokines IL-6, TNF-α, and KC (CXCL8) (Figure 6C). As such, we concluded that while Roflumilast is a used and FDA approved therapeutic, and influences human cilia, it may not work consistently or effectively in an in vivo mouse model of viral infection.

## Discussion

The large increase in pneumococcal burdens during coinfection would require significant resources for pneumococcal growth and replication. We saw that the EF3030 transcriptome reflects this necessity through the regulation of a diverse panel of genes, but mainly through its differential regulation of metabolic aspects. Our findings showed that upon encountering the human lung epithelium, multiple scavenging and nutrient processing genes are upregulated in order to grow. However, during pH1N1 coinfection, several of these are no longer as important, likely due to the effect of pH1N1 on the human lung epithelium.

Analysis of the reads mapping to the EF3030 transcriptome in our dataset showed a large increase in bacterial signal (∼30x) upon influenza infection (Figure 2B). Although this isn’t a conclusive measure of increased pneumococcal burdens upon pH1N1 infection, others have shown that bacterial CFUs increase significantly and remain elevated upon secondary pneumococcal infection (22). Despite increased EF3030 burdens in coinfection, host gene expression differences were largely incapable of distinguishing nHBEC cellular pathways unique to the coinfection state, implying that EF3030 may have little impact on IAV infected cells. This could be due to multiple reasons. Perhaps EF3030 being a non-invasive strain (18) means it is naturally less virulent, or 6 hours of incubation was not sufficient to perturb the IAV infected cells with 72 hours of accumulated toxins, or rather the nHBEC cells saw IAV as more of an existential threat, and the response to secondary EF3030 infection would take less precedence.

As for the relatively milder increase in pH1N1 reads mapped in our study (∼50%, Figure 2B), one study showed that viral titers initially increase then decline with time, typically after 7 days (22). This could be happening in our dataset as well, however 72+6 hours of post bacterial infection may not be sufficient to capture the eventual viral decline. Interestingly, in the coinfected samples, viral reads mapped were still much higher than those mapped to the pneumococcus (Supplemental Table 1), suggesting the virus was always an active participant in the environment. What’s more, viral titers during coinfection in vitro showed no changes in one study (23), so whether viral replication consistently increases may be very dependent on conditions and model systems, and therefore, remains inconclusive.

Global nutritional metabolism and regulation in the pneumococcus is maintained by CodY, and the catabolite control protein A (CcpA). CodY has been shown to regulate carbon metabolism, amino acid metabolism, and iron uptake (24). CcpA regulates several metabolic genes directly and indirectly, with some studies suggesting it controls ∼19% of all pneumococcal genes (25). Both studies have demonstrated that regulation by these two proteins has an impact on overall virulence of the pneumococcus. The complex level of regulation of these two master regulators has not yet been perfectly characterized. However, using our EF3030 transcriptome we have detailed the specific gene expression profiles for the regulated genes described in the RegPrecise database (Figure 4). We highlighted specific genes showing responses to one of the three conditions we studied, namely, EF3030 in vitro, Host+EF3030 & Host+EF3030+pH1N1. For example, in the CcpA regulon, carbohydrate sources such as mannose, galactose, maltose, galactosamine and sialic acid were upregulated specifically during EF3030 infection. Pyruvate usage, glycerol, maltodextrin and ATP synthetase genes were upregulated in the pH1N1 coinfection state. Similarly, in CodY, genes that trend toward upregulation in a pH1N1 coinfection state were expectedly amino acid sources such as glutamate, threonine, aspartate and other branched chain amino acids. While these global regulators offer a snapshot into parts of most pneumococcal metabolism, specific complete regulons are discussed below.

Regulons MalR and GlnR both seem to mimic complex patterns of regulation based on EF3030 infection or pH1N1 coinfection seen in CcpA and CodY. Both have some genes that were differentially regulated due to CcpA or CodY. However, it has been shown that CcpA performs some regulation over only some of the operons involved in maltose utilization (26). Specifically, the genes EF3030_RS01340 and EF3030_RS10400 through EF3030_RS10410 are regulated by CcpA and the rest of the regulon by MalR (26, 27). This is clearly evident from the Maltose utilization heatmap (Figure 4), and we can conclude that Maltose utilization occurs in both infection and pH1N1 coinfection, however different genes are involved. Similarly, in the case of GlnR, there is some discrepancy in the genes involved in the regulon. For example, the GlnR regulon in RegPrecise for Spn TIGR4, consists of 4 operons and 1 gene (Figure 3B, Figure 4). However, one study conducted in D39 established that only the genes *glnRA* (EF3030_RS02375 - EF3030_RS02380), *glnPQ/zwf* (EF3030_RS05240 - EF3030_RS05250) and *gdhA* (EF3030_RS06285) are part of the regulon and went on to show that a *ΔglnA* mutant and the *ΔglnP-glnA* double mutant (*ΔglnAP*) were attenuated in a mouse colonization model (28). Based on our infection condition specific expression pattern seen only for these genes (Figure 4), we conclude that the same is true in EF3030. The other 2 operons may be regulated in some strains such as TIGR4 but not necessarily all pneumococci. In line with our in vivo knock out mutant experiments, one study also showed severe attenuation of the *glnPQ/zwf* knockout (EF3030_RS05240 - EF3030_RS05250) in D39 in a pneumonia and septicemia mouse infection model (29).

We also saw regulation specific to EF3030 encountering the nHBEC cells in the form of zinc homeostasis (by AcdR) and transport proteins (by YwzG) (Figure 4). It has been established that the pneumococcus requires zinc to survive and responds to environmental Zn^2+^ levels (30). This study also established in D39 that there are two Zn^2+^ uptake mechanisms, one involving the AcdA gene, the other utilizing histidine triad proteins (Phts), and their functions are complementary (30, 31). Based on our expression data there is a slight difference between EF3030 infection and pH1N1 coinfection among the Phts. This is interesting as it was shown that Phts require surface attachment to function and provide zinc (30), supporting our results that EF3030 is in a biofilm state during infection but not pH1N1 coinfection, and that potentially different mechanisms of zinc acquisition are involved during pH1N1 coinfection. As for YwzG, relatively little is known in pneumococci. YwzG is classified as a PadR family regulator in *Bacillus subtilis*, as it controls the expression of phenolic acid decarboxylase, which detoxifies harmful phenolic acids. It is quite possible that the presence of toxic phenolic compounds can arise during nHBEC infection and pH1N1 coinfection. The upregulation of these genes in our data would suggest detoxification but this remains to be verified.

Multiple Spn regulons showed stark expression profiles specific to pH1N1 coinfection, namely fatty acid biosynthesis, riboflavin biosynthesis, pyrimidine metabolism and NADH energy metabolism (Rex) (Figure 4). The RegPrecise database asserts that pyrimidine metabolism and riboflavin biosynthesis in Spn TIGR4 are regulated (wholly or partially) through non-coding RNA mechanisms. Genes involved in riboflavin metabolism have been shown to induce activation of mucosal-associated invariant T cells (32), similar to our pH1N1 coinfection state. During *in vivo* infection, in the presence of host immune cells, this could lead to such T cell activation. This also supports our relatively quiescent EF3030 infection state from the host perspective as riboflavin genes are not as expressed (Figure 4). It has also been determined that deactivation of fatty acid biosynthesis results in a colony phase variation (33) and can also affect quorum sensing (34). It is quite possible that this allows EF3030 to leave its colonization state during pH1N1 coinfection and help turn invasive. Our group and others have also previously shown that fatty acid biosynthesis is downregulated not only in the blood (35-37) but also in human plasma (37). The EF3030+pH1N1 coinfection may also make EF3030 more adaptable to transitioning into the bloodstream. The PyrR regulon is fairly understudied in pneumococci apart from its identification and mechanism of regulation (38). The downregulation of pyrimidine genes we see during coinfection could be due to the pH1N1 nHBEC cell lysis providing an excess of free nucleotides for uptake. Lastly, while also understudied, the upregulated trend of EF3030 genes in our pH1N1 coinfection state for the Rex regulon suggests increased NADH usage (39) obtained potentially from nHBEC niacin released from pH1N1 lysed cells.

One of the most interesting and striking patterns of gene expression we observed was for lactose/galactose utilization, two forms of N-acetylglucosamine utilization, and sialic acid utilization, with all these being exclusive to just EF3030 infection (Figure 4). Utilization of these carbon sources is known to be favorable for biofilm formation (40), suggesting an EF3030 biofilm status and trend toward being in a colonization mode on nHBEC without external influences such as immune cells or coinfections. Downregulation of the LacR regulon was also observed in D39 at high glucose concentrations (41). This potentially adapts EF3030 transitioning into the glucose-rich bloodstream during pH1N1 coinfection. Another study by our group also showed that in such a pneumococcal state, the host response in a mouse nasopharynx model is also minimal (36). This is seen here in EF3030 on HBEC cells in both the EF3030 and host transcriptome. However, upon pH1N1 coinfection, EF3030 turns invasive and alters its metabolism to better suit the new environment. What’s more, all these carbon sources are acquired from epithelial host glycoconjugates (40). During pH1N1 coinfection, these are no longer used as nutritional sources, allowing EF3030 to turn invasive. Such strong high expression and tight regulation of these genes could also contribute to EF3030 being a preferentially lung-associated pneumococcal strain.

In the case of DE virulence genes in EF3030 (Figure 4), we see most genes downregulated in EF3030+pH1N1 coinfection relative to EF3030 infection. Exceptions to this were *ksgA, ppmA*, and *htrA*. Little is known about *ksgA* (EF3030_RS09680/SP_1992), with some studies suggest it is a cell wall surface anchor protein (42) and is a potential adhesin binding to collagen and lactoferrin (43). PpmA (EF3030_RS04725/SP_0981) is a lipoprotein involved in protease maturation (43). HtrA is a well-studied extracellular pneumococcal serine protease, and a recent study was performed with the EF3030 strain and its HtrA knockout mutants (44). It was shown to play a role in adherence to host cells and colonization, along with CbpG and PrtA (44). As we see from the virulence heatmap (Figure 4), during EF3030+pH1N1 coinfection, EF3030 prefers utilization of HtrA over the other two for adherence to nHBEC cells. As for those virulence genes important in just EF3030 infection, we have already discussed a few in their regulons such as neuraminidase genes, choline binding proteins (CbpD/CbpG), and histidine triad proteins (Phts), are all utilized in the absence of pH1N1 (Figure 4). Thus, EF3030 expression profiles we observed suggest that presence of influenza coinfection triggers a change away from a biofilm state, and away from feeding off host glycoconjugates. One study showed that introduction of influenza triggered a dispersion of pneumococcal biofilms in vitro (45). These researchers went on to identify ∼70 genes that were transcriptionally regulated specifically to induce dispersion from a pneumococcal biofilm state. Eighteen of these genes were found to follow the same trend in our Host+EF3030+pH1N1 vs. Host+EF3030 DE dataset. Yet another study demonstrated this in vivo in mice, where influenza coinfection and dispersion from a biofilm induced hypervirulence and invasive disease in pneumococcal strains EF3030 and D39 (46). Together with our data this suggests that the pneumococcus exists as a colonizing biofilm on the lung epithelium, but during influenza coinfection, a dispersion of this biofilm occurs, turning it into its invasive state, where it follows a different metabolic profile.

Some pneumococcal virulence factors in influenza superinfections were explored for essentiality in an in vivo CRISPRi-seq model, and showed that the pneumococcal capsule, together with *purA, bacA* and *pacL* genes were required for transmission, but interestingly not pneumolysin (7). We see in our dataset that in EF3030, most of the genes in the capsular locus (EF3030_RS01695-EF3030_RS01745) are not DE passed cutoffs when it encounters the host lung epithelium, but their expression is elevated and identified in our black WGCNA module (Figure 3A). We also saw that pneumolysin (EF3030_RS09370) was significantly downregulated when encountering the host lung epithelium, but *bacA* (EF3030_RS02235) had no changes in expression across all conditions. All the genes found relevant during influenza superinfection in CRISPRi-seq, except for *bacA*, a bacitracin susceptibility gene (likely due to it being identified in vivo in mice which could potentially contain other lung microbes), follow suit with our dataset in terms of expression trends. This combined with known dispersion from biofilm and increased bacterial loads observed in our study and others, support the possibility that EF3030 switches toward a transmission phenotype during influenza coinfection.

Platt et al. have demonstrated using transcriptomics and proteomics that pneumococcal metabolism can be altered by IAV alone, not requiring the presence of the host (47). Certain aspects of Spn metabolism can be explained by direct interaction with IAV. For example, using proteomics, they identified a strong downregulation in pyrimidine metabolism in the presence of IAV, regardless of the host. We observed this same response in our EF3030 transcriptomics (Figure 4). However, while they saw galactose metabolism through LacR upregulated, we see it is transcriptionally downregulated (Figure 4). Interestingly, when they compared the transcriptomes of Spn + IAV (15 min p.i.) to Spn alone, they found the GlnRA operon was downregulated. Again, we see the inverse in Spn EF3030 in our study. It is important to note that they performed most of their experiments in TIGR4, a highly invasive strain. These could be strain specific phenotypes that are relevant to the different diseases they can cause (EF3030-Lung adapted vs. TIGR4-invasive). Interestingly, in our in vivo KO experiments of TIGR4, the GlnRA KO was less attenuated in the coinfection state, fitting their TIGR4 observation, but EF3030 may show a different phenotype. However, the GlnRA downregulation was absent by the 30 min time point in their study (47).

As we established an invasive pneumococcal state upon influenza coinfection versus colonization during infection, we investigated if influenza infection in the host resulted in differential expression of genes that aid invasive disease. For example, two known pneumococcal host surface receptors PECAM-1 and RSPA (48, 49), that are used by bacteria to achieve host cell invasion, were upregulated about ∼30x (∼3-5 Log_2_ Fold Change, Supplemental Table 4) upon influenza infection in our study, potentially aiding the invasive pneumococcus.

Finally, several pathways identified by GO analyses mediate upregulation of diverse immune regulatory systems (Supplemental Table 4) to counteract infection, these observations are consistent with other studies showing the involvement of dendritic cell induced cytokine signaling and IL-17 signaling (11, 50). These data suggest that pH1N1 infection and EF3030+pH1N1 coinfection of human lung epithelial cells result in the upregulation of cytokines and signaling to recruit host immune cells. On the other end of the spectrum, we observed that several downregulated genes (>2500 genes) are related to cilial beating and microtubule-based cell movement in our GO analysis, suggesting that this is a significant biological process in response to influenza infection, with and without EF3030. This phenotype was also observed in our pilot experiments which followed identical protocols of infection, where a marked decrease in cilia was observed with immunofluorescence (Supplemental Figure 1). Such cilial regulation is consistent with prior influenza and pneumococcal infection studies (51), also involving upregulation of IL-17 and IFN genes seen in our data. As the host response to the influenza virus is overwhelming, we intend to delve into more specific aspects of human regulatory pathways that could be involved such as transport, protein channels etc., in a later study.

## Methods

### Isolation and culture of differentiated human bronchial epithelial cells and sample generation

Primary human bronchial epithelial cells were obtained from human lung tissue through the tissue donation program by the International Institute for the Advancement of Medicine using a previously published method (52). Cells were harvested from the human tracheobronchial tissues and plated onto collagen-coated 100-mm plates (Biocoat; Becton Dickinson), in BEGM medium to ∼80% confluency, followed by passaging onto 6.5-mm Transwell inserts (0.4-μm pore size) with FNC coating (0407, AthenaES) and differentiated at an Air-Liquid Interface (ALI), as previously described (53), except with the modification that cells were grown and expanded on collagen coated plates (Advanced Biomatrix 5028-10EA) in complete growth media from LifeLine Cell Technology (LL-0023) and subsequently differentiated to air with complete Pneumacult ALI media (05001) from Stemcell Technologies. Cells were incubated for approximately two weeks until appearance of ciliated cells and mucus production was seen (53). Following culture of differentiated human bronchial epithelial cells on transwells, cells were infected with pH1N1 or mock infected (no pH1N1 controls) for 72 h (∼50% of cells infected with IAV). pH1N1 infected cells were then exposed to 1000 CFUs of Streptococcus pneumoniae strain EF3030 for 6 h. For pneumococcal EF3030 control samples, ∼50,000 CFUs of EF3030 were cultured in ALI media for 6 hours. 1ml of RNA protect was then added to all transwells and then stored in 5ml of RNAprotect Bacteria Reagent in 50 ml falcon tubes at -80C.

### RNA isolation, library construction, sequencing, and transcriptomics data analyses

On the day of RNA isolation, transwell samples were thawed and the membranes cut out using a scalpel. RNAprotect and isolated membranes were centrifuged at 10,000 rpm, to pellet the membrane and any dislodged cells, and the supernatant discarded. Pellets were then incubated in 100 μL of lysis buffer (10 μL of mutanolysin, 20 μL of proteinase K, 30 μL of lysozyme, 40 μL of TE buffer) for 10 minutes. Followed by mechanical disruption in 600 μL RLT buffer (RNeasy Mini Kit, Qiagen) containing 1% ß-mercaptoethanol, using a motorized pestle for 30 seconds (36). RNA was then captured on the RNeasy Mini Kit columns with DNase treatment on column (Qiagen protocol).

Extracted RNA was quantitated using a Bioanalyzer. Ribosomal RNA was depleted using the RiboZero rRNA Removal Kits for Gram-positive bacteria and/or for human/mouse/rat (Illumina). 300 bp-insert RNA-seq Illumina libraries were constructed using ∼1.0 μg of enriched mRNA that was fragmented then used for synthesis of strand-specific cDNA using the NEBnext Ultra Directional RNA Library Prep Kit (NEB-E7420L). The cDNA was purified between enzymatic reactions and the size selection of the library performed with AMPure SpriSelect Beads (Beckman Coulter Genomics). The titer and size of the libraries was assessed on the LabChip GX (Perkin Elmer) and with the Library Quantification Kit (Kapa Biosciences). RNA-seq was conducted on 150 nt paired-end runs of the Illumina NovaSeq 6000 platform using two or three biological replicates for each condition (36).

FASTQ files were mapped to their respective genomes using HISAT (54) for Homo sapiens and Bowtie (55) for pneumococcal EF3030 and viral pH1N1 genomes. Gene expression counts for all samples were then estimated using HTseq (56). The counts tables were then used for analyses and estimation of differentially expressed (DE) genes. Rarefaction curves were generated from counts data. Principal Component Analyses (PCAs) and dendrograms were generated in R based on normalized Variance Stabilized Transformation (VST) counts acquired using the DESeq2 R package (12). For human DE gene estimation, infected samples were compared to their respective uninfected control samples as the baseline using DEseq2 and filtered using an FDR cutoff of ≤0.05 and an absolute Log_2_ Fold Change cutoff of ≥1. For bacterial DE gene estimation, each EF3030 infected sample was compared to EF3030 grown in ALI media as the baseline, using DEseq2 and filtered using an FDR cutoff of ≥0.05 and an absolute Log2 Fold Change cutoff of ≥1. Common and unique DE genes for both species were determined using Upset plots (R package UpsetR (57)) and individual heatmaps of DE genes were generated based on Z-scores of VST counts (R package DESeq2). WGCNA (13) was performed on DEseq2 low expression filtered EF3030 genes (1924 genes). Modules were then merged using the default merge eigengene dissimilarity threshold of 0.25. Four modules demonstrated gene expression patterns fitting the sample condition metadata. These were tabulated to 835 EF3030 genes that represent the major changes across the EF3030 transcriptome in this study. All fastq files and associated metadata are uploaded to the Gene Expression Omnibus (GEO) repository (see Data availability).

### Orthology

Orthologous genes between EF3030 and TIGR4 were determined using PanOCT v3.23 (58) with the parameters -S Y -M Y -H Y -F 1.33 -c 0,25,50,75,100 -T.

### Bacterial gene regulon analysis

Bacterial DE gene lists were used to determine DE regulons using the online RegPrecise (15) database for *Streptococcus pneumoniae* TIGR4, with the condition that at least 2 genes of the regulon were differentially expressed. Bacterial WGCNA module amino acid sequences were uploaded to the eggNOG-mapper (14) online tool for COG classification.

### Gene Ontology (GO) analysis

GO ontologies and associated figures were estimated for all Human DE genes that from all comparisons (FDR ≤ 0.05 and an absolute Log_2_ Fold Change of ≥1) using Cluster Profiler version 4 (17).

### Animals

Male and female 8-week-old C57BL/6 mice were purchased from Jackson Laboratory (Bar Harbor, ME). Mice were housed at 21°C in a 12-hour light/dark cycle and given food and water ad libitum. Experimental procedures were approved by the Institutional Animal Care and Use Committee at The University of Alabama at Birmingham (Protocol #21579).

### Knock-out mutants

Mutants were created by allelic exchange in strain TIGR4 using the EF3030-TIGR4 orthologous genes. Mutagenic PCR constructs were generated by amplifying the upstream and downstream DNA fragments flanking the gene(s) of interest followed by fusion of these fragments with the Janus Cassette using a HiFi assembly master mix (NEB). Transformation of TIGR4 with the mutagenic construct (100ng/ml) was performed by natural transformation which was induced using competence-stimulating peptide variant 2 (CSP-2) as previously described (36). Blood agar plates supplemented with kanamycin (300 mg/L) were used for selection.

### Infection and Therapeutic Intervention

Mice were anesthetized with 2.5% isoflurane and intranasally infected with 500 plaque-forming units (PFU) of influenza A virus PR8 (H1N1 A/Puerto Rico/8/1934) in 50 μl. On day 3 post-infection, mice were treated with 1 mg/kg Roflumilast (Sigma #SMG 1099) or vehicle (PBS/0.02% Tween 80/0.5% hydroxycellulose) in 100 μl via oral gavage. Briefly, a 1.5-inch 20 gauge feeding needle (Med Vet International) attached to a 1 mL syringe was gently inserted down the esophagus into the stomach before injection of the dose. Mice were not anesthetized during this procedure and were observed after treatment for signs of discomfort or poor recovery. Treatment was given every 24 hours for 5 days during which the mice were also weighed.

### Cytokine Measurement in Lung Homogenate

Mice were sacrificed using isoflurane and whole lungs were harvested and rinsed in PBS before being homogenized in 1 mL of PBS. Homogenates were spun down at 5000 g for 5 minutes and supernatants collected. Cytokine levels for IL-6, TNF-α, and CXCL1/KC were measured using R&D DuoSet ELISA kits (IL-6 #DY406, TNF-α #DY410, CXCL1/KC #DY453) according to the manufacturer’s instructions.

### Detection of H1N1

A plaque-forming assay was used to assess the amount of H1N1 in the lung homogenates of infected mice. Madin-Darby canine kidney (MDCK) cells were seeded in a 12-well cell culture dish at a density of 1 × 10^5^ cells/well and grown to confluence. Lung homogenate dilutions (10^−1^-10^−10^) were prepared in inoculating media (1x serum-free Eagle’s Minimum Essential Medium and Trypsin TPCK) and 500 μl added per well. Homogenates were incubated for 1 hour then removed. 1 mL of overlay media (2x serum-free EMEM and 2.4% Avicell) was added to each well and the plate was further incubated for 7 days (long enough for plaque-formation). The overlay media was removed, and the cells fixed with 10% formalin overnight at 4°C. Fixative was removed, and the cells were washed with deionized water. The cells were stained with 0.05% neutral red for 1 hour at room temperature, then rinsed again with DI water. The plates were allowed to dry completely before plaques enumerated. Virus titer was determined by [#plaques/(volume of inoculum x dilution factor)].

### Statistical Analysis

Multiple comparisons were performed by one-way analysis of variance with Tukey’s post-hoc test. All results shown are standard error of the mean (SEM). Each individual point represents an individual animal sample. Asterisks denote the level of significance observed: * = p ≤ 0.05; **** = p ≤ 0.0001. Statistical analyses were calculated using GraphPad Prism 9 software (La Jolla, CA).

## Data Availability

R code/scripts used to generate figures and data presented are available on GitHub at https://github.com/admelloGithub/ExVivoSpnH1N1_v2. The data reported in this paper are in the process of being deposited in the Gene Expression Omnibus (GEO) database, https://www.ncbi.nlm.nih.gov/geo (accession no. GSEXXXXXX).

## Author contributions

HT, CJO and KSH designed the study; AD performed experiments and major data analyses; JRL, JLT, EM, and HNR performed experiments and analyzed data; All authors contributed to manuscript writing.

## Acknowledgements

This project was funded in part by federal funds from the National Institute of Allergy and Infectious Diseases, National Institutes of Health, Department of Health and Human Services under grant number U19AI110820, and R01AI156893. We also acknowledge essential support provided by members of the Institute for Genome Sciences’ Maryland Genomics core, the Genome Informatics Core, and High-Performance Computing Core.

## Supplemental Figure Legends

**Supplemental Figure 1.**
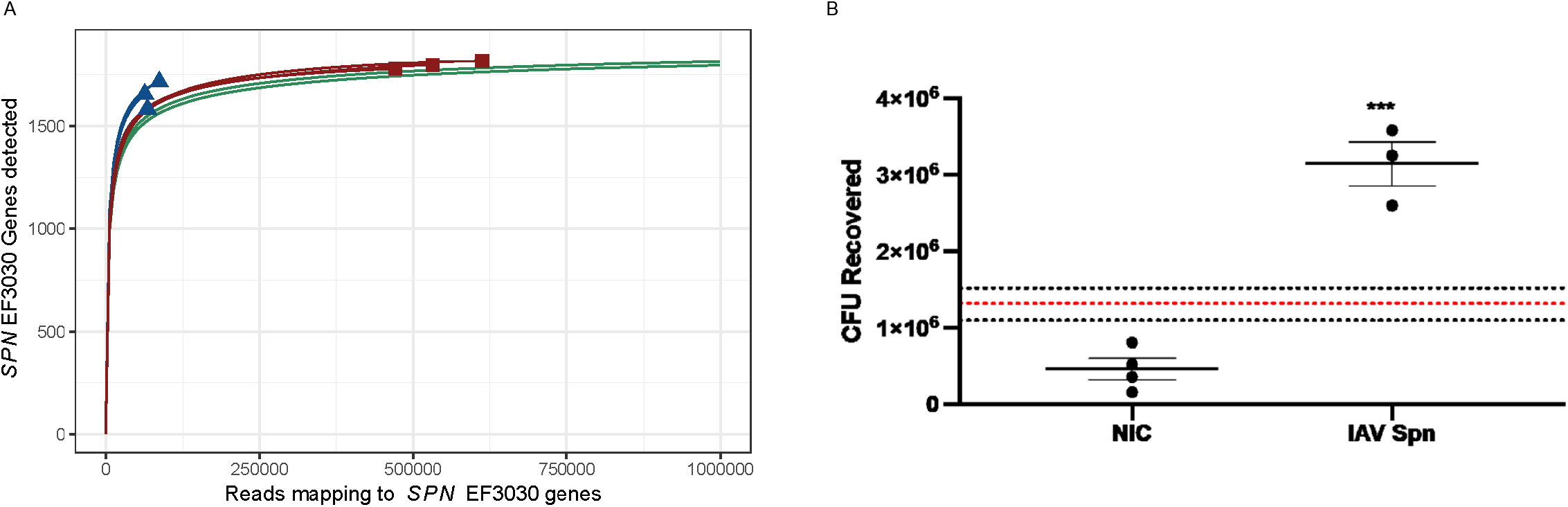
Rarefaction curves of RNA-seq reads mapped to the EF3030 genome and EF3030 colony forming units (CFU). A) Rarefaction curves of EF3030 samples. Samples whose curves plateau indicate sufficient sequencing depth for the majority of EF3030 genes analyzed. B) EF3030 CFU counts for EF3030 monoinfection (no-influenza control, NIC) vs. influenza A virus + EF3030 coinfected nHBEC cells (IAV Spn).

**Supplemental Figure 2.**
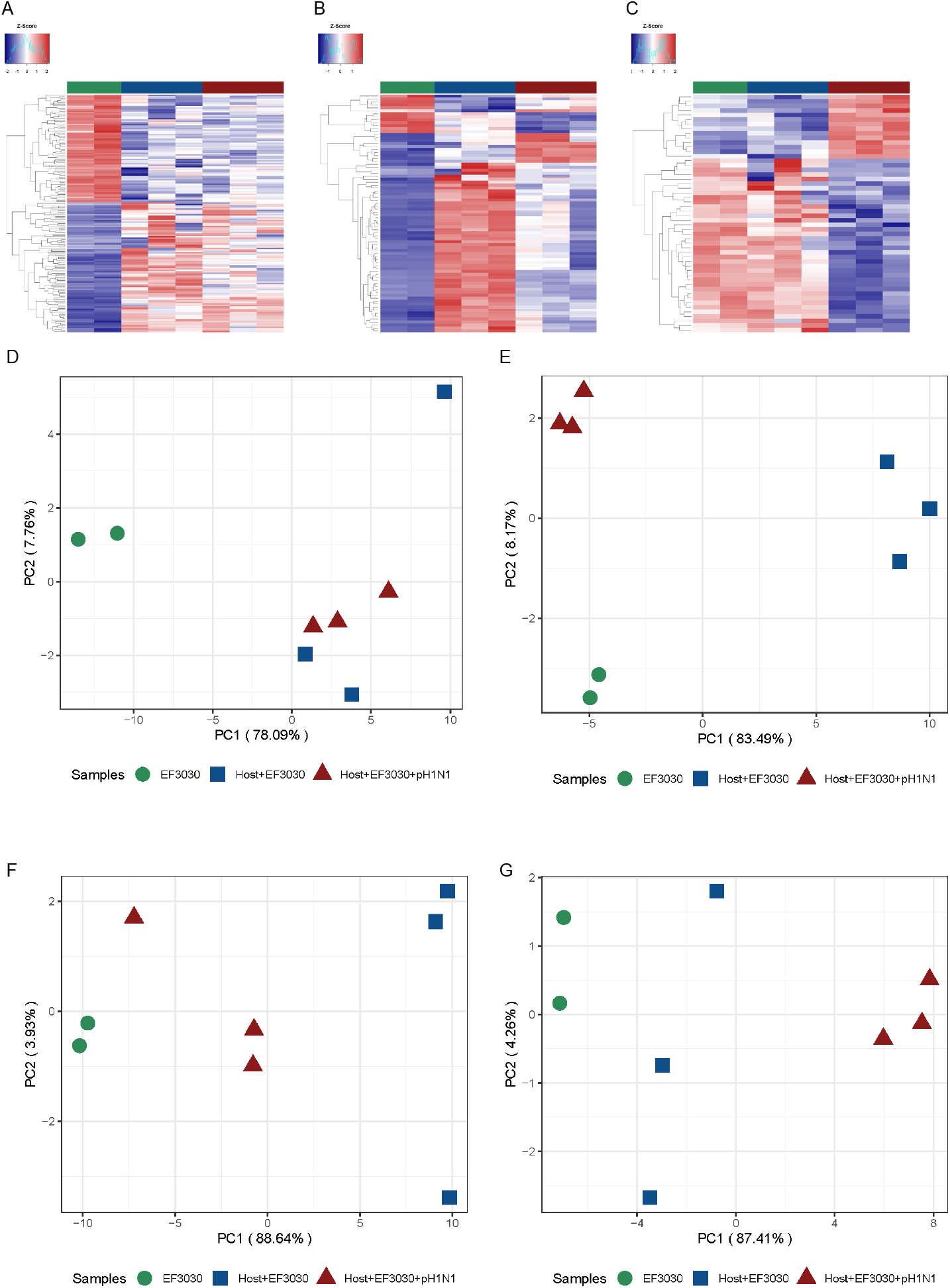
Z-scored Heatmaps of differentially expressed (DE) EF3030 gene intersects and principal component analyses (PCA) of EF3030 WGCNA modules in Figure 3A. A) Heatmap of 163 DE genes shared in comparisons of Host+EF3030+pH1N1 vs. EF3030 and Host+EF3030 vs. EF3030, showing genes influenced by nHBEC. B) Heatmap of 80 DE genes shared in all comparisons, showing genes having unique condition-specific levels of expression. C) Heatmap of 52 DE genes shared in comparisons of Host+EF3030+pH1N1 vs. EF3030 and Host+EF3030+pH1N1 vs Host+EF3030, showing genes influenced by pH1N1. Entire DE gene lists are shown in Supplemental Table 3. D-G) PCA of each of the EF3030 WGCNA modules in Figure 3A, black, blue, green, and pink, respectively. Sample clustering and high proportions of variation seen in PC1 (78-88%) suggest the genes in these modules effectively capture condition-specific expression.

**Supplemental Figure 3.**
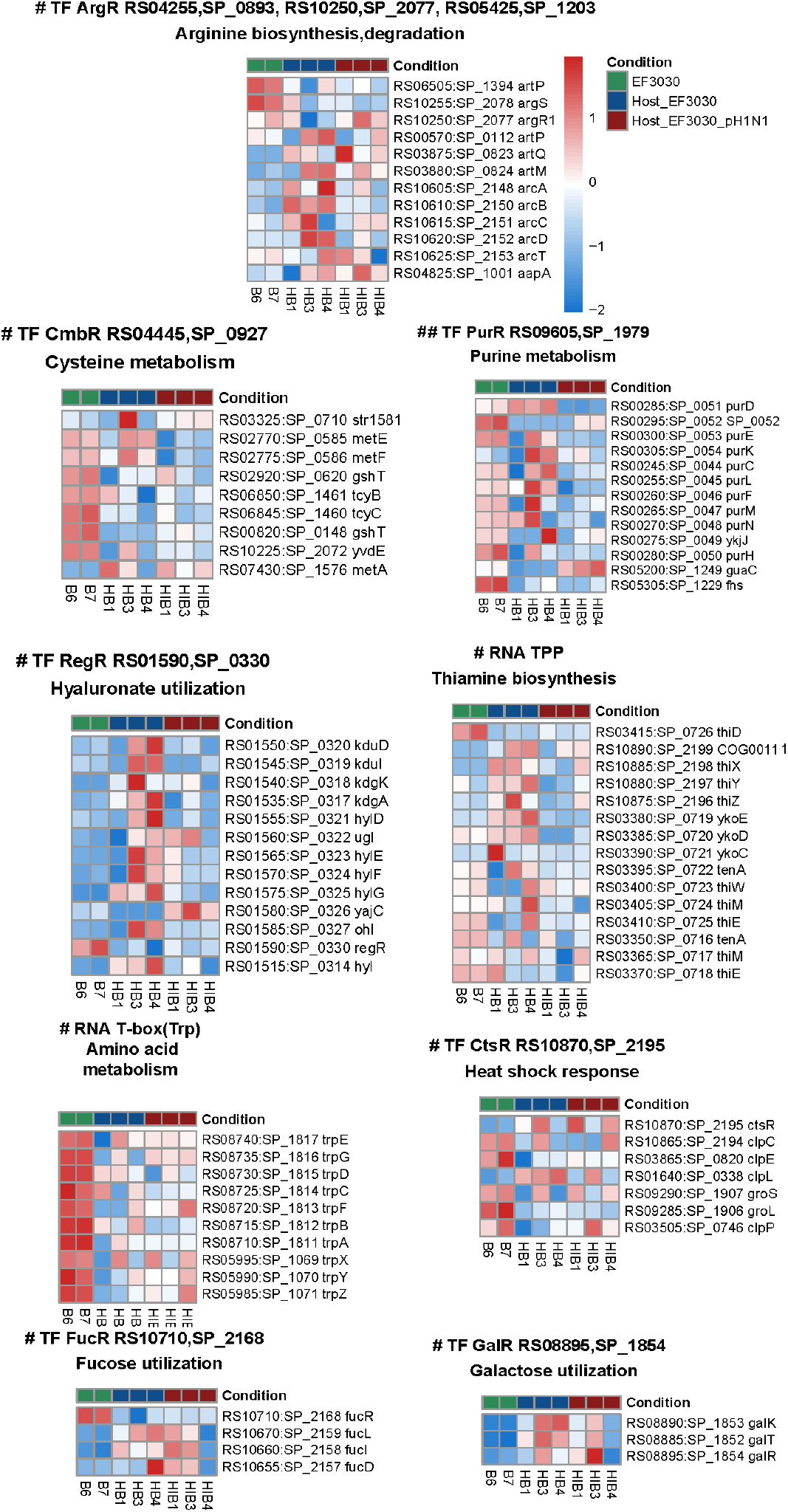
Z-scored Heatmaps of Regprecise EF3030 regulons having at least 2 genes identified as differentially expressed. Shown are the 9 regulons identified as differentially regulated which had no consistent condition-specific gene expression, and therefore were not shown in Figure 4.

**Supplemental Figure 4.**
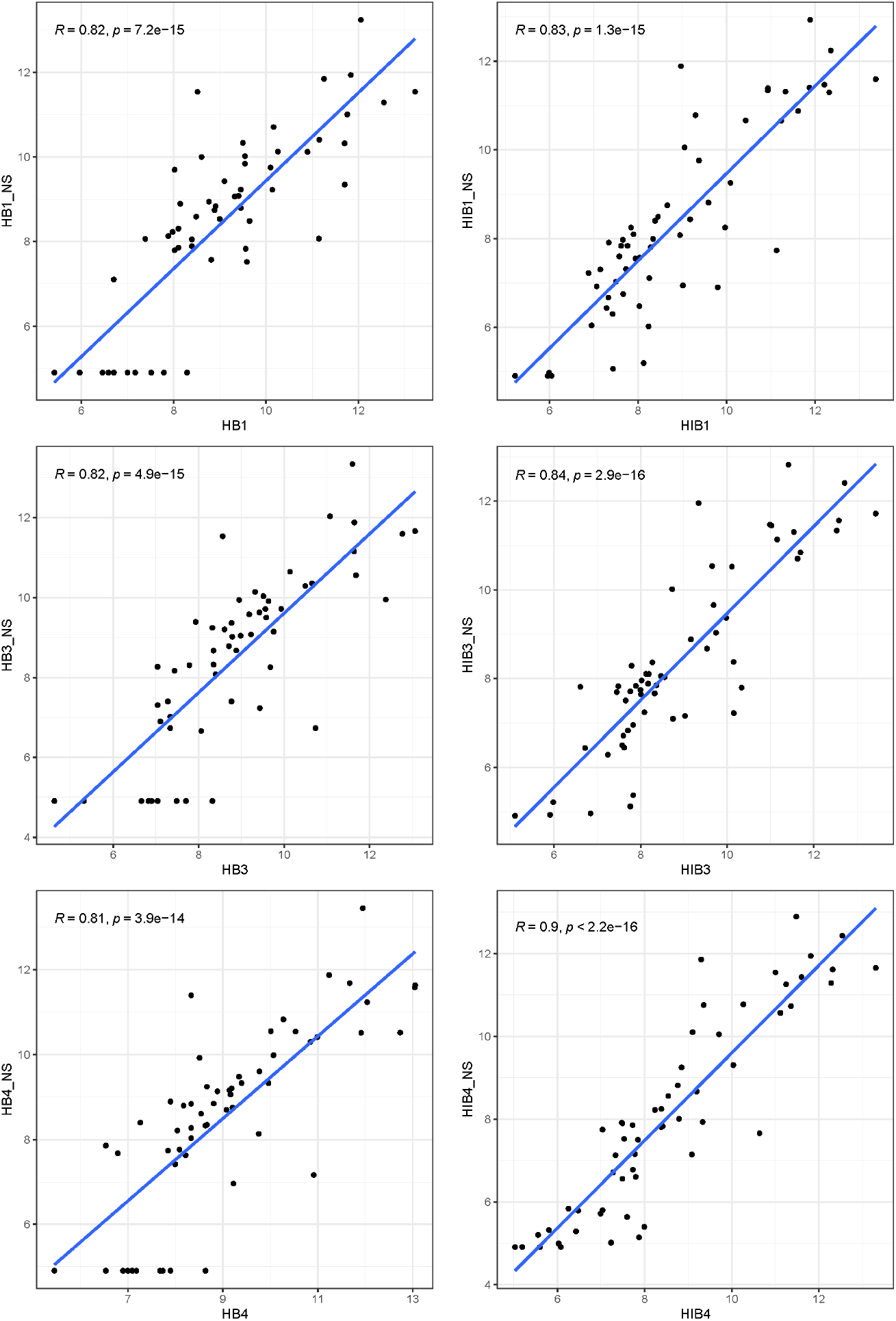
Spearman correlations of RNA-Seq vs. Nanostring for 63 EF3030 genes. X axes are DESeq2 Variance Stabilized Transformation (VST) counts. Y axes are normalized Nanostring expression values. Both axes are on a Log_2_ scale.

**Supplemental Figure 5.**
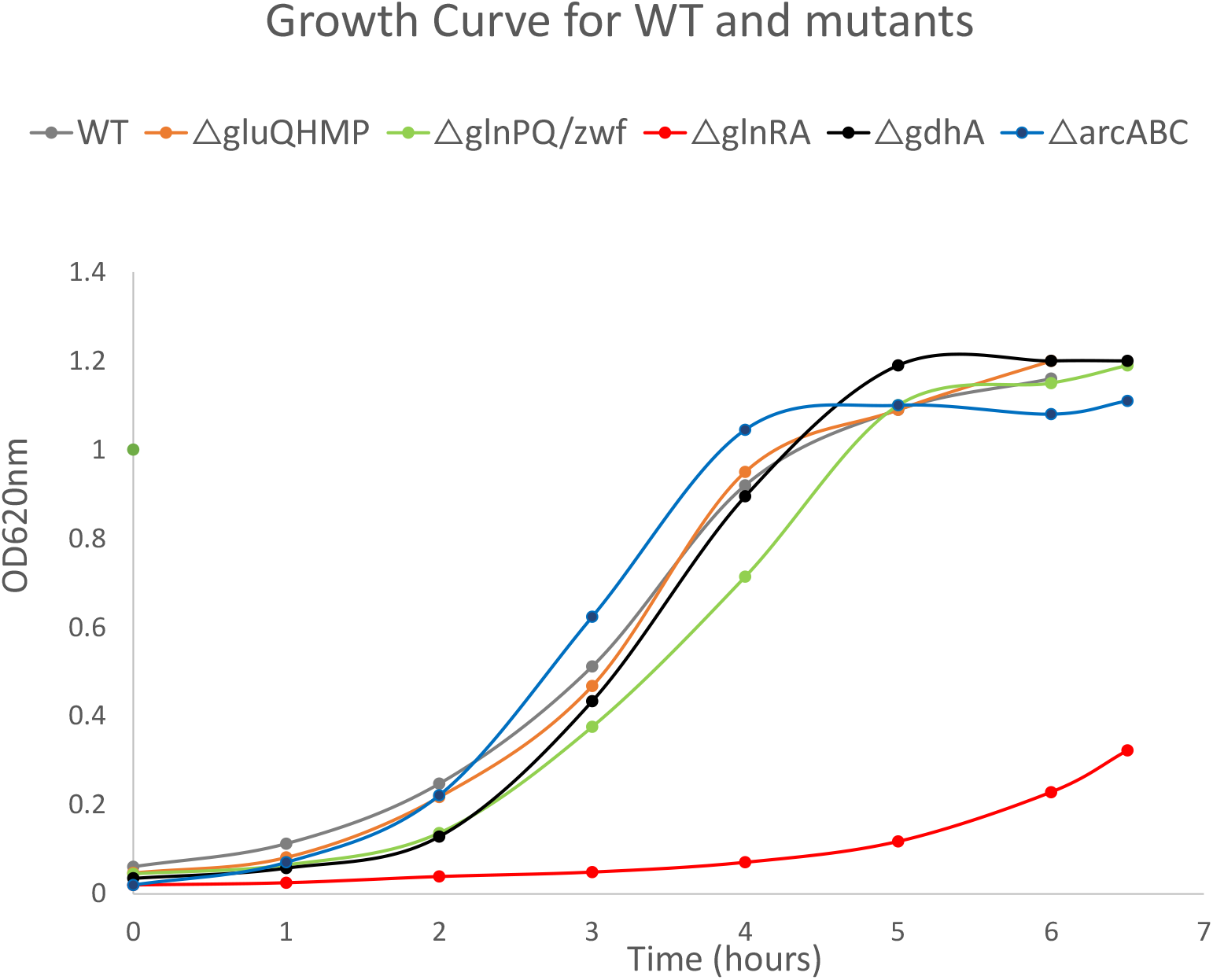
TIGR4 wildtype vs. TIGR4 knockout GlnR regulon mutant growth curves. Optical density measurements of *Streptococcus pneumoniae* TIGR4 wt/mutant growth over 6-7 hours in liquid media.

## References

1. Klein EY, Monteforte B, Gupta A, Jiang W, May L, Hsieh YH, et al. The frequency of influenza and bacterial coinfection: a systematic review and meta-analysis. Influenza Other Respir Viruses. 2016;10(5):394–403.

2. MacIntyre CR, Chughtai AA, Barnes M, Ridda I, Seale H, Toms R, et al. The role of pneumonia and secondary bacterial infection in fatal and serious outcomes of pandemic influenza a(H1N1)pdm09. BMC Infect Dis. 2018;18(1):637.

3. Short KR, Habets MN, Hermans PW, Diavatopoulos DA. Interactions between Streptococcus pneumoniae and influenza virus: a mutually beneficial relationship? Future Microbiol. 2012;7(5):609–24.

4. Morens DM, Taubenberger JK, Fauci AS. Predominant role of bacterial pneumonia as a cause of death in pandemic influenza: implications for pandemic influenza preparedness. J Infect Dis. 2008;198(7):962–70.

5. Diavatopoulos DA, Short KR, Price JT, Wilksch JJ, Brown LE, Briles DE, et al. Influenza A virus facilitates Streptococcus pneumoniae transmission and disease. FASEB J. 2010;24(6):1789–98.

6. Rowe HM, Livingston B, Margolis E, Davis A, Meliopoulos VA, Echlin H, et al. Respiratory Bacteria Stabilize and Promote Airborne Transmission of Influenza A Virus. mSystems. 2020;5(5).

7. Liu X, Kimmey JM, Matarazzo L, de Bakker V, Van Maele L, Sirard JC, et al. Exploration of Bacterial Bottlenecks and Streptococcus pneumoniae Pathogenesis by CRISPRi-Seq. Cell Host Microbe. 2021;29(1):107–20 e6.

8. Rowe HM, Karlsson E, Echlin H, Chang TC, Wang L, van Opijnen T, et al. Bacterial Factors Required for Transmission of Streptococcus pneumoniae in Mammalian Hosts. Cell Host Microbe. 2019;25(6):884–91 e6.

9. Reinoso-Vizcaino NM, Cian MB, Cortes PR, Olivero NB, Hernandez-Morfa M, Pinas GE, et al. The pneumococcal two-component system SirRH is linked to enhanced intracellular survival of Streptococcus pneumoniae in influenza-infected pulmonary cells. PLoS Pathog. 2020;16(8):e1008761.

10. Pettigrew MM, Marks LR, Kong Y, Gent JF, Roche-Hakansson H, Hakansson AP. Dynamic changes in the Streptococcus pneumoniae transcriptome during transition from biofilm formation to invasive disease upon influenza A virus infection. Infect Immun. 2014;82(11):4607–19.

11. Ambigapathy G, Schmit T, Mathur RK, Nookala S, Bahri S, Pirofski LA, et al. Double-Edged Role of Interleukin 17A in Streptococcus pneumoniae Pathogenesis During Influenza Virus Coinfection. J Infect Dis. 2019;220(5):902–12.

12. Love MI, Huber W, Anders S. Moderated estimation of fold change and dispersion for RNA-seq data with DESeq2. Genome Biol. 2014;15(12):550.

13. Langfelder P, Horvath S. WGCNA: an R package for weighted correlation network analysis. BMC Bioinformatics. 2008;9:559.

14. Cantalapiedra CP, Hernandez-Plaza A, Letunic I, Bork P, Huerta-Cepas J. eggNOG-mapper v2: Functional Annotation, Orthology Assignments, and Domain Prediction at the Metagenomic Scale. Mol Biol Evol. 2021;38(12):5825–9.

15. Novichkov PS, Kazakov AE, Ravcheev DA, Leyn SA, Kovaleva GY, Sutormin RA, et al. RegPrecise 3.0--a resource for genome-scale exploration of transcriptional regulation in bacteria. BMC Genomics. 2013;14:745.

16. Gamez G, Castro A, Gomez-Mejia A, Gallego M, Bedoya A, Camargo M, et al. The variome of pneumococcal virulence factors and regulators. BMC Genomics. 2018;19(1):10.

17. Wu T, Hu E, Xu S, Chen M, Guo P, Dai Z, et al. clusterProfiler 4.0: A universal enrichment tool for interpreting omics data. Innovation (Camb). 2021;2(3):100141.

18. Junges R, Maienschein-Cline M, Morrison DA, Petersen FC. Complete Genome Sequence of Streptococcus pneumoniae Serotype 19F Strain EF3030. Microbiol Resour Announc. 2019;8(19).

19. Joskova M, Mokry J, Franova S. Respiratory Cilia as a Therapeutic Target of Phosphodiesterase Inhibitors. Front Pharmacol. 2020;11:609.

20. Rabe KF. Update on roflumilast, a phosphodiesterase 4 inhibitor for the treatment of chronic obstructive pulmonary disease. Br J Pharmacol. 2011;163(1):53–67.

21. Mata M, Martinez I, Melero JA, Tenor H, Cortijo J. Roflumilast inhibits respiratory syncytial virus infection in human differentiated bronchial epithelial cells. PLoS One. 2013;8(7):e69670.

22. Smith AM, Adler FR, Ribeiro RM, Gutenkunst RN, McAuley JL, McCullers JA, et al. Kinetics of coinfection with influenza A virus and Streptococcus pneumoniae. PLoS Pathog. 2013;9(3):e1003238.

23. Ouyang K, Woodiga SA, Dwivedi V, Buckwalter CM, Singh AK, Binjawadagi B, et al. Pretreatment of epithelial cells with live Streptococcus pneumoniae has no detectable effect on influenza A virus replication in vitro. PLoS One. 2014;9(3):e90066.

24. Hendriksen WT, Bootsma HJ, Estevao S, Hoogenboezem T, de Jong A, de Groot R, et al. CodY of Streptococcus pneumoniae: link between nutritional gene regulation and colonization. J Bacteriol. 2008;190(2):590–601.

25. Carvalho SM, Kloosterman TG, Kuipers OP, Neves AR. CcpA ensures optimal metabolic fitness of Streptococcus pneumoniae. PLoS One. 2011;6(10):e26707.

26. Afzal M, Shafeeq S, Manzoor I, Kuipers OP. Maltose-Dependent Transcriptional Regulation of the mal Regulon by MalR in Streptococcus pneumoniae. PLoS One. 2015;10(6):e0127579.

27. Puyet A, Ibanez AM, Espinosa M. Characterization of the Streptococcus pneumoniae maltosaccharide regulator MalR, a member of the LacI-GalR family of repressors displaying distinctive genetic features. J Biol Chem. 1993;268(34):25402–8.

28. Hendriksen WT, Kloosterman TG, Bootsma HJ, Estevao S, de Groot R, Kuipers OP, et al. Site-specific contributions of glutamine-dependent regulator GlnR and GlnR-regulated genes to virulence of Streptococcus pneumoniae. Infect Immun. 2008;76(3):1230–8.

29. Hartel T, Klein M, Koedel U, Rohde M, Petruschka L, Hammerschmidt S. Impact of glutamine transporters on pneumococcal fitness under infection-related conditions. Infect Immun. 2011;79(1):44–58.

30. Plumptre CD, Hughes CE, Harvey RM, Eijkelkamp BA, McDevitt CA, Paton JC. Overlapping functionality of the Pht proteins in zinc homeostasis of Streptococcus pneumoniae. Infect Immun. 2014;82(10):4315–24.

31. Plumptre CD, Eijkelkamp BA, Morey JR, Behr F, Counago RM, Ogunniyi AD, et al. AdcA and AdcAII employ distinct zinc acquisition mechanisms and contribute additively to zinc homeostasis in Streptococcus pneumoniae. Mol Microbiol. 2014;91(4):834–51.

32. Hartmann N, McMurtrey C, Sorensen ML, Huber ME, Kurapova R, Coleman FT, et al. Riboflavin Metabolism Variation among Clinical Isolates of Streptococcus pneumoniae Results in Differential Activation of Mucosal-associated Invariant T Cells. Am J Respir Cell Mol Biol. 2018;58(6):767–76.

33. Zhang J, Ye W, Wu K, Xiao S, Zheng Y, Shu Z, et al. Inactivation of Transcriptional Regulator FabT Influences Colony Phase Variation of Streptococcus pneumoniae. mBio. 2021;12(4):e0130421.

34. Aggarwal SD, Gullett JM, Fedder T, Safi JPF, Rock CO, Hiller NL. Competence-Associated Peptide BriC Alters Fatty Acid Biosynthesis in Streptococcus pneumoniae. mSphere. 2021;6(3):e0014521.

35. Shenoy AT, Brissac T, Gilley RP, Kumar N, Wang Y, Gonzalez-Juarbe N, et al. Streptococcus pneumoniae in the heart subvert the host response through biofilm-mediated resident macrophage killing. PLoS Pathog. 2017;13(8):e1006582.

36. D’Mello A, Riegler AN, Martinez E, Beno SM, Ricketts TD, Foxman EF, et al. An in vivo atlas of host-pathogen transcriptomes during Streptococcus pneumoniae colonization and disease. Proc Natl Acad Sci U S A. 2020;117(52):33507–18.

37. Pettersen JS, Hog FF, Nielsen FD, Moller-Jensen J, Jorgensen MG. Global transcriptional responses of pneumococcus to human blood components and cerebrospinal fluid. Front Microbiol. 2022;13:1060583.

38. Aprianto R, Slager J, Holsappel S, Veening JW. High-resolution analysis of the pneumococcal transcriptome under a wide range of infection-relevant conditions. Nucleic Acids Res. 2018;46(19):9990–10006.

39. Afzal M, Shafeeq S, Kuipers OP. NADH-Mediated Gene Expression in Streptococcus pneumoniae and Role of Rex as a Transcriptional Repressor of the Rex-Regulon. Front Microbiol. 2018;9:1300.

40. Blanchette KA, Shenoy AT, Milner J, 2nd, Gilley RP, McClure E, Hinojosa CA, et al. Neuraminidase A-Exposed Galactose Promotes Streptococcus pneumoniae Biofilm Formation during Colonization. Infect Immun. 2016;84(10):2922–32.

41. Suo W, Guo X, Zhang X, Xiao S, Wang S, Yin Y, et al. Glucose levels affect MgaSpn regulation on the virulence and adaptability of Streptococcus pneumoniae. Microb Pathog. 2022;174:105896.

42. van de Garde MDB, van Westen E, Poelen MCM, Rots NY, van Els C. Prediction and Validation of Immunogenic Domains of Pneumococcal Proteins Recognized by Human CD4(+) T Cells. Infect Immun. 2019;87(6).

43. Jahn K, Kohler TP, Swiatek LS, Wiebe S, Hammerschmidt S. Platelets, Bacterial Adhesins and the Pneumococcus. Cells. 2022;11(7).

44. Ali MQ, Kohler TP, Burchhardt G, Wust A, Henck N, Bolsmann R, et al. Extracellular Pneumococcal Serine Proteases Affect Nasopharyngeal Colonization. Front Cell Infect Microbiol. 2020;10:613467.

45. Chao Y, Marks LR, Pettigrew MM, Hakansson AP. Streptococcus pneumoniae biofilm formation and dispersion during colonization and disease. Front Cell Infect Microbiol. 2014;4:194.

46. Marks LR, Davidson BA, Knight PR, Hakansson AP. Interkingdom signaling induces Streptococcus pneumoniae biofilm dispersion and transition from asymptomatic colonization to disease. mBio. 2013;4(4).

47. Platt MP, Lin YH, Penix T, Wiscovitch-Russo R, Vashee I, Mares CA, et al. A multiomics analysis of direct interkingdom dynamics between influenza A virus and Streptococcus pneumoniae uncovers host-independent changes to bacterial virulence fitness. PLoS Pathog. 2022;18(12):e1011020.

48. Orihuela CJ, Mahdavi J, Thornton J, Mann B, Wooldridge KG, Abouseada N, et al. Laminin receptor initiates bacterial contact with the blood brain barrier in experimental meningitis models. J Clin Invest. 2009;119(6):1638–46.

49. Iovino F, Molema G, Bijlsma JJ. Platelet endothelial cell adhesion molecule-1, a putative receptor for the adhesion of Streptococcus pneumoniae to the vascular endothelium of the blood-brain barrier. Infect Immun. 2014;82(9):3555–66.

50. Kuri T, Sorensen AS, Thomas S, Karlsson Hedestam GB, Normark S, Henriques-Normark B, et al. Influenza A virus-mediated priming enhances cytokine secretion by human dendritic cells infected with Streptococcus pneumoniae. Cell Microbiol. 2013;15(8):1385–400.

51. LeMessurier KS, Tiwary M, Morin NP, Samarasinghe AE. Respiratory Barrier as a Safeguard and Regulator of Defense Against Influenza A Virus and Streptococcus pneumoniae. Front Immunol. 2020;11:3.

52. Fulcher ML, Gabriel S, Burns KA, Yankaskas JR, Randell SH. Well-differentiated human airway epithelial cell cultures. Methods Mol Med. 2005;107:183–206.

53. Brand JD, Lazrak A, Trombley JE, Shei RJ, Adewale AT, Tipper JL, et al. Influenza-mediated reduction of lung epithelial ion channel activity leads to dysregulated pulmonary fluid homeostasis. JCI Insight. 2018;3(20).

54. Kim D, Langmead B, Salzberg SL. HISAT: a fast spliced aligner with low memory requirements. Nat Methods. 2015;12(4):357–60.

55. Langmead B, Trapnell C, Pop M, Salzberg SL. Ultrafast and memory-efficient alignment of short DNA sequences to the human genome. Genome Biol. 2009;10(3):R25.

56. Anders S, Pyl PT, Huber W. HTSeq--a Python framework to work with high-throughput sequencing data. Bioinformatics. 2015;31(2):166–9.

57. Conway JR, Lex A, Gehlenborg N. UpSetR: an R package for the visualization of intersecting sets and their properties. Bioinformatics. 2017;33(18):2938–40.

58. Fouts DE, Brinkac L, Beck E, Inman J, Sutton G. PanOCT: automated clustering of orthologs using conserved gene neighborhood for pan-genomic analysis of bacterial strains and closely related species. Nucleic Acids Res. 2012;40(22):e172.

